# *In situ* cell condensation-based cartilage tissue engineering via immediately implantable high-density stem cell core and rapidly degradable shell microgels

**DOI:** 10.1101/2024.04.20.590385

**Authors:** Sang Jin Lee, Oju Jeon, Yu Bin Lee, Daniel S. Alt, Aixiang Ding, Rui Tang, Eben Alsberg

## Abstract

Formation of chondromimetic human mesenchymal stem cells (hMSCs) condensations typically required *in vitro* culture in defined environments. In addition, extended *in vitro* culture in differentiation media over several weeks is usually necessary prior to implantation, which is costly, time consuming and delays clinical treatment. Here, this study reports on immediately implantable core/shell microgels with a high-density hMSC-laden core and rapidly degradable hydrogel shell. The hMSCs in the core formed cell condensates within 12 hours and the oxidized and methacrylated alginate (OMA) hydrogel shells were completely degraded within 3 days, enabling spontaneous and precipitous fusion of adjacent condensed aggregates. By delivering transforming growth factor-β1 (TGF-β1) within the core, the fused condensates were chondrogenically differentiated and formed cartilage microtissues. Importantly, these hMSC-laden core/shell microgels, fabricated without any *in vitro* culture, were subcutaneously implanted into mice and shown to form cartilage tissue via cellular condensations in the core after 3 weeks. This innovative approach to form cell condensations *in situ* without *in vitro* culture that can fuse together with each other and with host tissue and be matured into new tissue with incorporated bioactive signals, allows for immediate implantation and may be a platform strategy for cartilage regeneration and other tissue engineering applications.

## 1. Introduction

Articular cartilage defects frequently occur due to sporting injuries and other acute trauma, resulting in degeneration of the cartilage tissue.^[1]^ Due to low cellularity, lack of vasculature, and deficient production of the extracellular matrix, articular cartilage possesses poor self-regeneration capacity.^[2]^ Currently, micro-fracture is considered the standard of care for early stage, limited cartilage defect repair, but often fibrous cartilaginous scar tissue is formed with inferior mechanical properties.^[3]^ Mosaicplasty, in which small osteochondral cylinders from low weight bearing surfaces of the affected joint or another joint of the patient are transferred into the damaged site, suffers from donor site morbidity, limited availability, and varied long-term outcomes.^[4]^ An alternative treatment, autologous chondrocyte implantation, has inconsistent outcomes^[5]^ and requires multiple surgeries that make it a challenging task for the orthopaedic surgeon.^[6]^ Other strategies include implantation of decellularized allografts and synthetic biomaterials, which can result in poor integration with host tissue.^[7]^

Over the past several decades, cell-laden biomaterial scaffolding-based tissue engineering approaches have been widely explored in efforts to treat cartilage defects.^[8]^ Unfortunately, the use of biomaterial scaffolds to delivery cells for cartilage tissue engineering has faced several challenges, such as the interference of the biomaterial with critical cell-cell interactions, unsynchronized scaffold degradation and new tissue formation rates, tissue inhomogeneity, a low density of seeded cells, and potential immunogenicity of the scaffold biomaterials and their degradation by-products.^[9]^

To overcome these limitations of scaffold-based approaches, scaffold-free strategies to regenerate cartilage have been widely pursued where cells form condensations through cadherin mediated cell-cell interactions.^[10]^ Transforming growth factor-β1 (TGF-β1)-releasing microparticles have been incorporated into self-assembled human mesenchymal stem cell (hMSC) condensations to induce homogenous chondrogenic differentiation of hMSCs via more uniform and sustained chondroinductive factor presentation and to permit more rapid clinical implementation without extensive *in vitro* culture.^[10–11]^ However, even with the incorporation of TGF-β1-releasing microparticles within the cell condensation constructs, to date, their formation has still required *in vitro* culture in geometrically confined microenvironments prior to implantation.^[12]^ For example, cell condensations have been formed using microwell arrays,^[13]^ V-shape or round-bottom wells,^[14]^ the hanging drop method,^[15]^ Transwell^®^ membranes,^[16]^ hydrogel molds,^[17]^ and a 3D bioprinting slurry bath.^[9a]^ All these methods require *in vitro* culture for days to several weeks for cell-cell contacts and resulting condensations to form along with subsequent tissue maturation. This production process is labor-intensive, time-consuming, expensive, and delays implantation of such constructs.

To address the aforementioned issues, this study reports the first strategy, to our knowledge, that enables *in situ* cell condensation-based cartilage formation without *in vitro* culture. Immediately implantable high-density hMSC-laden core/shell microgels containing chondroinductive gelatin microparticles (GMs) were fabricated using a triaxial nozzle equipped with syringe pumps and airflow (Figure 1 and Figure S1). The core/shell microgels consisted of high-density hMSCs and TGF-β1-releasing GMs in the core and a thin rapidly degrading hydrogel shell layer of ionically crosslinked oxidized and methacrylated alginate (OMA). Unlike other reported core/shell systems,^[18]^ the *in situ* hydrolytically degradable OMA shell layer provided a temporary physical boundary for the hMSCs, enabling *in situ* cell condensation formation with incorporated TGF-β1-laden GMs. After rapid degradation of the OMA shell layer within a couple of days, resulting cell condensates could mature to form neocartilagenous tissues and fuse and integrate with neighboring condensates. Importantly, it was demonstrated that the immediately implantable core/shell microgels could form hMSC condensations that developed into cartilage tissue and fused with each other and host tissue in a subcutaneous mouse model without any prior *in vitro* culture.

**Figure 1.**
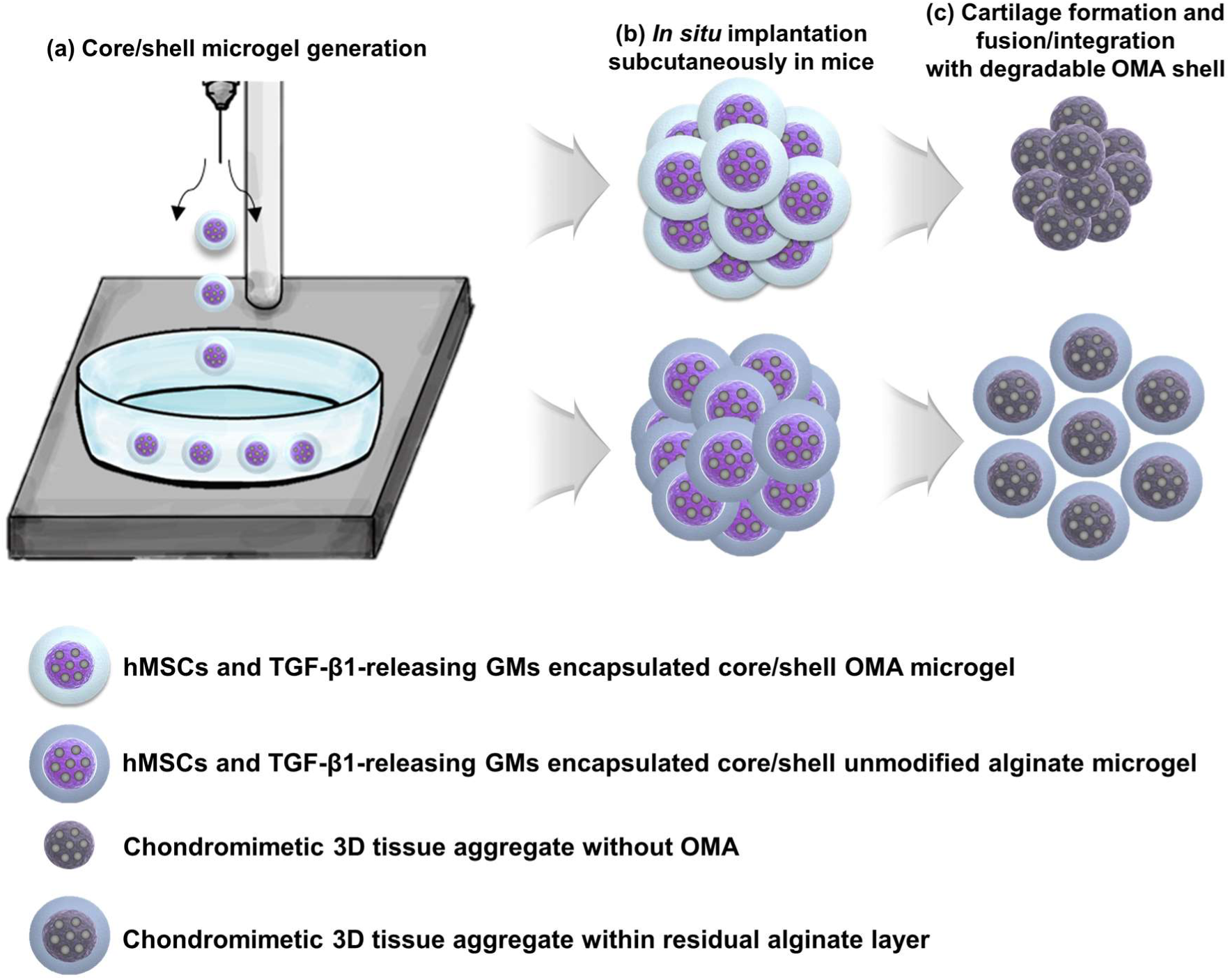
Schematic illustration of high-density hMSC and TGF-β1-releasing GM encapsulated core/shell OMA microgel. (a) Generation of core/shell microgels by triaxial needle system, (b) *in situ* subcutaneous implantation in mice without *in vitro* culture, and (c) hMSC condensation formation and subsequent chondrogenesis with neotissue fusion and integration with each other and host tissue in mice. Due to OMA’s ability to degrade, the OMA hydrogel shell layer goes away following hMSC condensation formation and the encapsulated TGF-β1-releasing GMs simultaneously stimulate hMSCs to undergo chondrogenesis. Finally, cartilage matrix forms and fuses with host tissue without a preceding *in vitro* culture period. In contrast, the unmodified alginate shell layer degrades very slowly, inhibiting neotissue fusion and integration.

## 2. Results

### 2.1 Synthesis of OMA and preparation of core/shell microgels

Unique to this core/shell system reported here is development of a shell material that (1) is stable over the course of a couple of days to enable in situ condensation formation in the core and (2) then degrades rapidly to enable fusion of the cell condensations with each other and host tissue. To serve this purpose, OMA was employed, which is dually crosslinkable by divalent cations and application of low level UV light. Increasing the degree of oxidation makes the alginate uronate residues more susceptible to hydrolysis and increases the degradation rate of the hydrogels.^[19]^ The actual oxidation degrees of oxidized alginates were 1.55 (2 % theoretical) and 3.36 % (4 % theoretical) (Figure S2 and Table S1), and the actual methacrylation degrees of the OMAs formed with 2 and 4 % theoretical oxidation at 40% theoretical methacrylation were 9.61 (2-OMA) and 10.51 % (4-OMA), respectively (Figure S3 and Table S1).

To form spherical core/shell hydrogel constructs, a triaxial nozzle equipped with syringe pumps and air-flow was utilized. A detailed schematic illustration of the high-density cell-laden core/shell microgel fabrication method is depicted in Figure 1 and Figure S1. Cells only or cells with TGF-β1 loaded GMs are extruded in the center and an unmodified alginate or OMA macromer solution is extruded surrounding the cells. Air-flow, the most exterior layer of the triaxial configuration, interupts the inner streams, resulting in a spherical core/shell droplet shape. These particles are released into and crosslinked in a bath containing a CaCl_2_ solution. In this manner, core/shell microgels could be continuously produced in a high-throughput manner (Figure S Movie 1).

The degradation kinetics of PBS core/ OMA shell microgels was then characterized. Bright-field images of cell-free core/2-/4-OMA shell hydrogel particles were obtained for 48 hours in media under cell culture conditions. The OMA core/shell microgels exhibited a spherical shape at 0h (Figure 2a). As the OMA shell degraded, the spherical structure of the OMA core/shell microgels gradually began to degrade over time, and most of the hydrogel shell was gone by 48 h. Biochemical quantification demonstrated that the 4-OMA shell degradation rate was faster than the 2-OMA (Figure 2b). The outer OMA shell also degraded rapidly within 48-72 h when cells were incorporated into the core (Figure. S4a and b), corroborating the OMA degradation findings with the cell-free core/shell particles.

**Figure 2.**
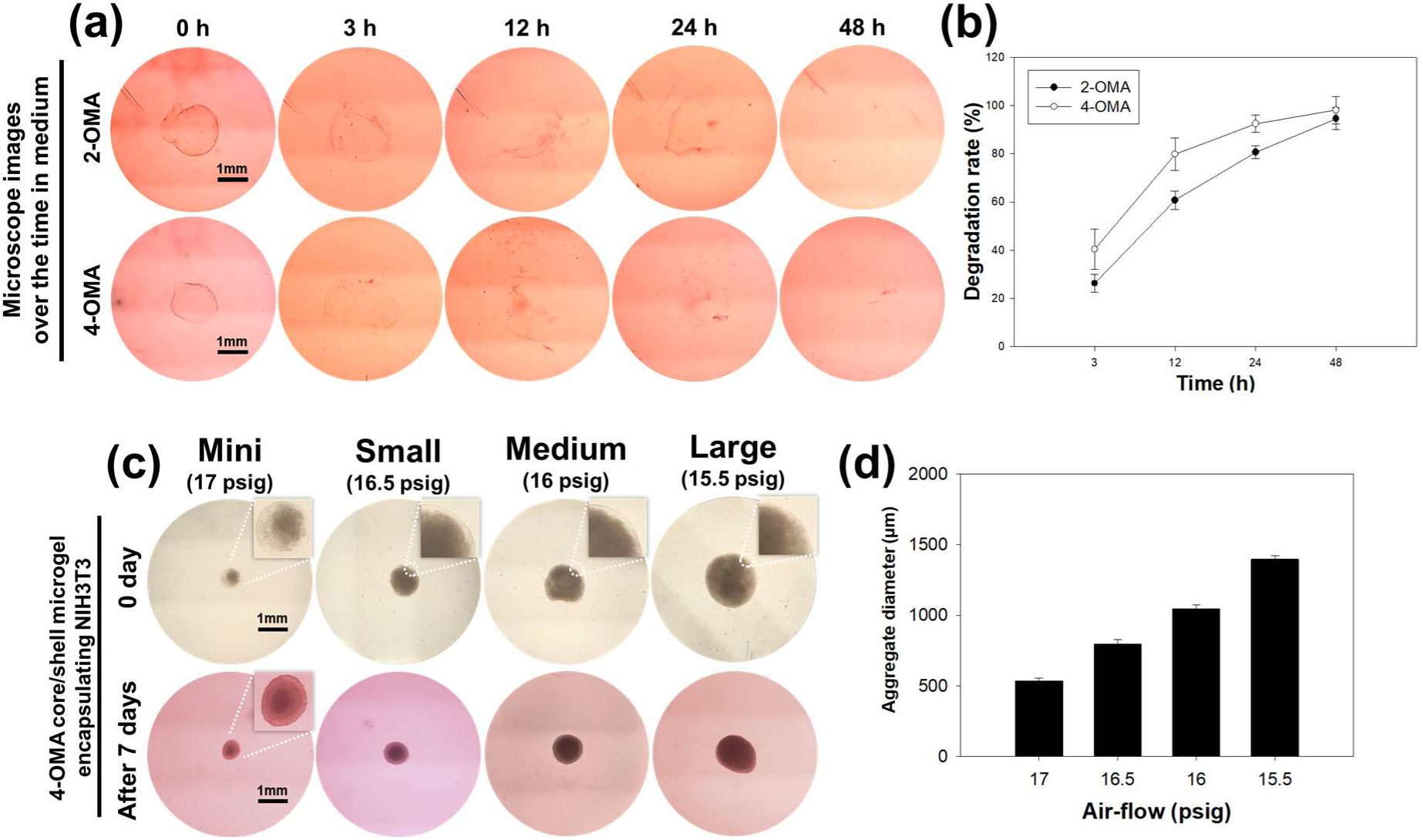
*In vitro* degradation kinetics of cell-free core / 2-/4-OMA shell microgels over 48 h. Each microgel was prepared without cells in the core filled by PBS. (a) The resultant microgels exhibited a round particle shape at 0 h, but this spheroidal structure gradually began to collapse over the time. The OMA components were nearly gone within 48 h in both groups. (b) Quantification of the degraded amount of 2-/4-OMA was measured by DMMB assay. The degradation rate of 4-OMA was faster than 2-OMA due to its higher oxidation degree. **Generation of different sizes of 4-OMA core/shell microgels.** (c) Different sizes of NIH3T3 encapsulated core/ 4-OMA shell microgels were produced by modulating the air-flow. After 7 days of culture, various sizes of spheroidal NIH3T3 aggregates were obtained and (d) their average diameters were measured.

Core/shell droplet composition and size can be modulated by varying the rate of the OMA macromer solution extrusion, cell extrusion and air-flow. For example, fabricated high-density cell-laden core/shell microgel size could be controlled by changing the air flow rate of the outer nozzle of the triaxial nozzle apparatus (Figure 2c, d, and S4c). After 7 days of culture, the alginate component of the core/shell microgels was degraded and high-density cell condensations remained. The morphology (Figure S4d) and size (Figure S4e) of 3D cell spheroids formed via the traditional V-shape well plate method were similar to those of the cell condensations formed by the core/shell microgel after 7 days of cultivation.

### 2.2 Morphology change of the high-density cell-laden core/shell microgels and their fusion

To demonstrate the impact of shell degradation, core/shell microgels were generated using NIH3T3 cells and unmodified alginate, 2-OMA and 4-OMA. While the hydrogel shell layer in the unmodified alginate group remained after 21 days of culture, the hydrogel shell layers of both 2 and 4-OMA groups disappeared within several days (Figure 3a and S5a). Interestingly, over time, the cell condensations self-assembled into a more spherical shape, likely due to cell-cell adhesions and cell cytoskeletal contractile forces. When the cell-laden core/shell microgels were cultured in a well plate for 3 days, adjacent cell aggregates fused together after the OMA shell degraded (Figure 3b). These fused constructs maintained their shape while exposed to strong pipetting of PBS (Figure S Movie 2). Unmodified alginate did not allow cell aggregate fusion because residual alginate physically prevented contact between the cellular component of adjacent constructs. Thus, individual core/shell microgels remained in the PBS when the shell could not degrade. When the cell condensations were cultured for 21 days, no difference in DNA amount was observed between two OMA groups (Figure S5b). In addition, high cell viability was observed at 7 days (Figure S5c).

**Figure 3.**
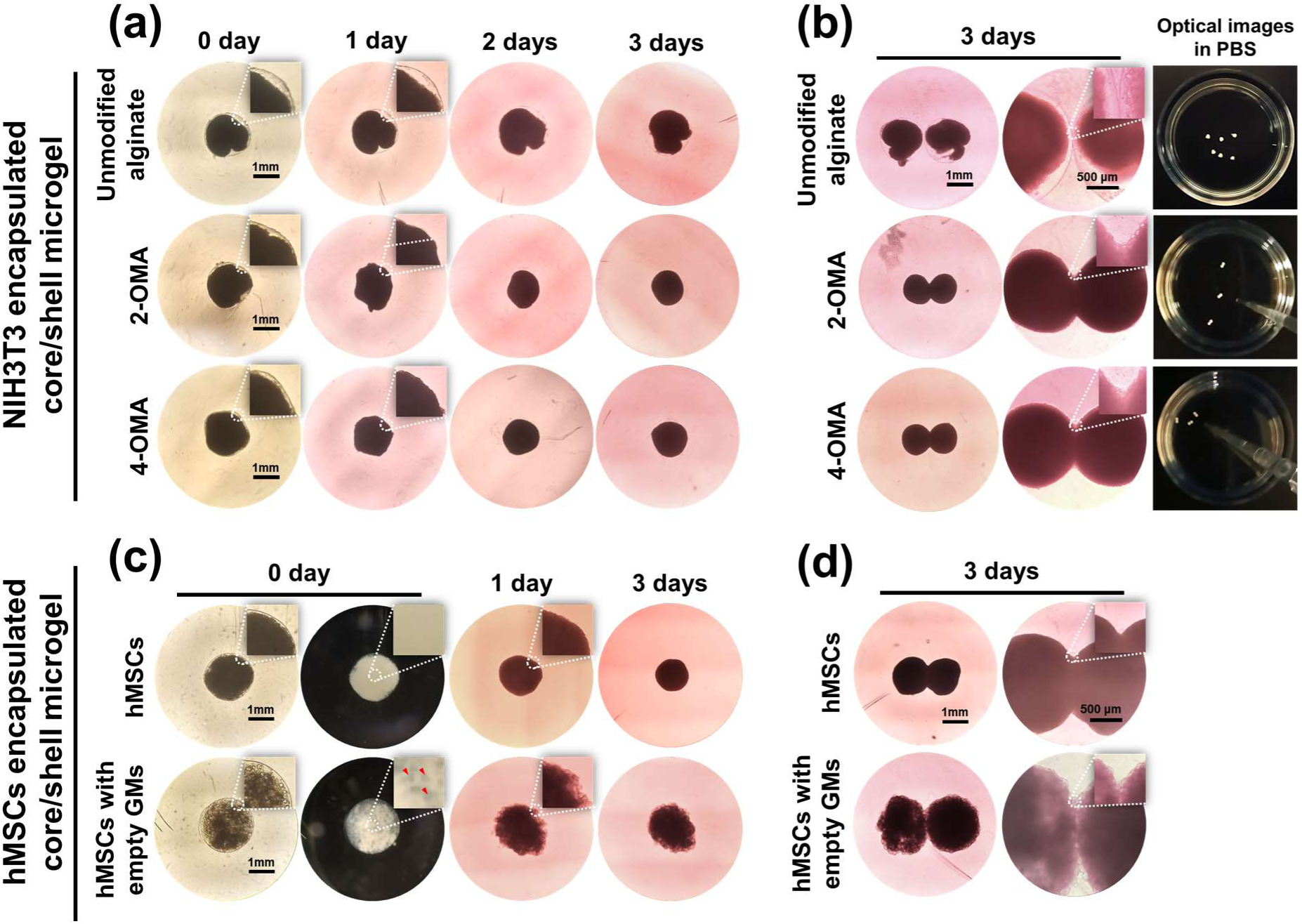
Morphology change of NIH3T3 encapsulated single and paired core/shell microgels over 3 days. (a) In the unmodified alginate group, the hydrogel shell layer continuously retained the NIH3T3s. In both the 2 and 4-OMA groups, a round spherically shaped cell condensation was gradually formed over time as the hydrogel shell layer degraded. Small inset images display the presence and absence of the hydrogel shell layer over 3 days. (b) Paired NIH3T3 encapsulated core/shell microgels were cultivated together in the same well plate for 3 days. 2-OMA and 4-OMA groups allowed for cell aggregate fusion after shell degradation. Small inset images show the direct cellular coupling between paired aggregates. In contrast, the unmodified alginate group did not allow for aggregate fusion. The fused structures retained their structure when put them in PBS with strong pipetting in both OMA groups. However, the unmodified alginate group spread out individually in PBS. **Single and paired hMSC and hMSC/GMs core/ 4-OMA shell microgels.** (c) hMSC and hMSC/GM encapsulated 4-OMA groups both maintained stable cell condensations, similar to NIH3T3 encapsulated 4-OMA microgels. The GMs were uniformly distributed within the cell aggregates. (d) Paired hMSC and hMSC/GMs core/ 4-OMA shell microgels fused spontaneously upon shell degradation (small inset).

Next, hMSCs alone or with empty GMs were encapsulated in core/shell microgels made of 4-OMA (Figure 3c). Optical photomicrographs showed hMSC condensations after 1 day with and without GMs. In groups with incorporated GMs, the GMs were retained in the cell aggregates for 21 days (Figure S6). The size of hMSCs aggregates with GMs was larger than hMSCs aggregate without GMs due to the additional volume of the incorporated GMs. Adjacent cell aggregates spontaneously fused after the OMA shell degraded (Figure 3d).

### 2.3 Chondrogenic differentiation and analysis of core/shell microgels with encapsulated hMSCs *in vitro*

The capacity to drive the chondrogenic differentiation of the core-formed cell condensation via exogenous supplementation or incorporated GM-mediated local delivery of potent chondrogenic cytokine TGF-β1 was then investigated. Detailed *in vitro* experimental groups are described in Table S2 and Figure S7. After 2 and 3 weeks of *in vitro* culture, the level of GAG was significantly higher at 3 weeks than 2 weeks in all groups and DNA content was similar at both weeks within each group (Figure 4a). However, DNA content of groups 4 and 6 was significantly greater than groups 3 and group 5, respectively, because of double aggregates in groups 4 and 6. Almost no GAG was measured in group 1 due to absence of TGF-β1, and GAG and GAG/DNA was significantly greater in group 2 due to the exogenously supplied TGF-β1.^[20]^ When comparing groups 2 and 3, GAG and GAG/DNA contents of group 3 was higher than group 2 because incorporation of GMs could enhance chondrogenesis, as previously shown in non-core/shell hMSC aggregates.^[20]^ When comparing group 3 to 5 or group 4 to 6, content of GAG/DNA in groups 5 and 6 was significantly higher than in groups 3 and 4, respectively, at 3 weeks, indicating that localized and sustained delivery of TGF-β1 from GMs within condensations enhanced chondrogenesis in this system.^[11a]^ The comparison between single and double microgels (group 3 vs 4 and group 5 vs 6) also showed significantly greater GAG content in double aggregates than in single aggregates at 3 weeks.

**Figure 4.**
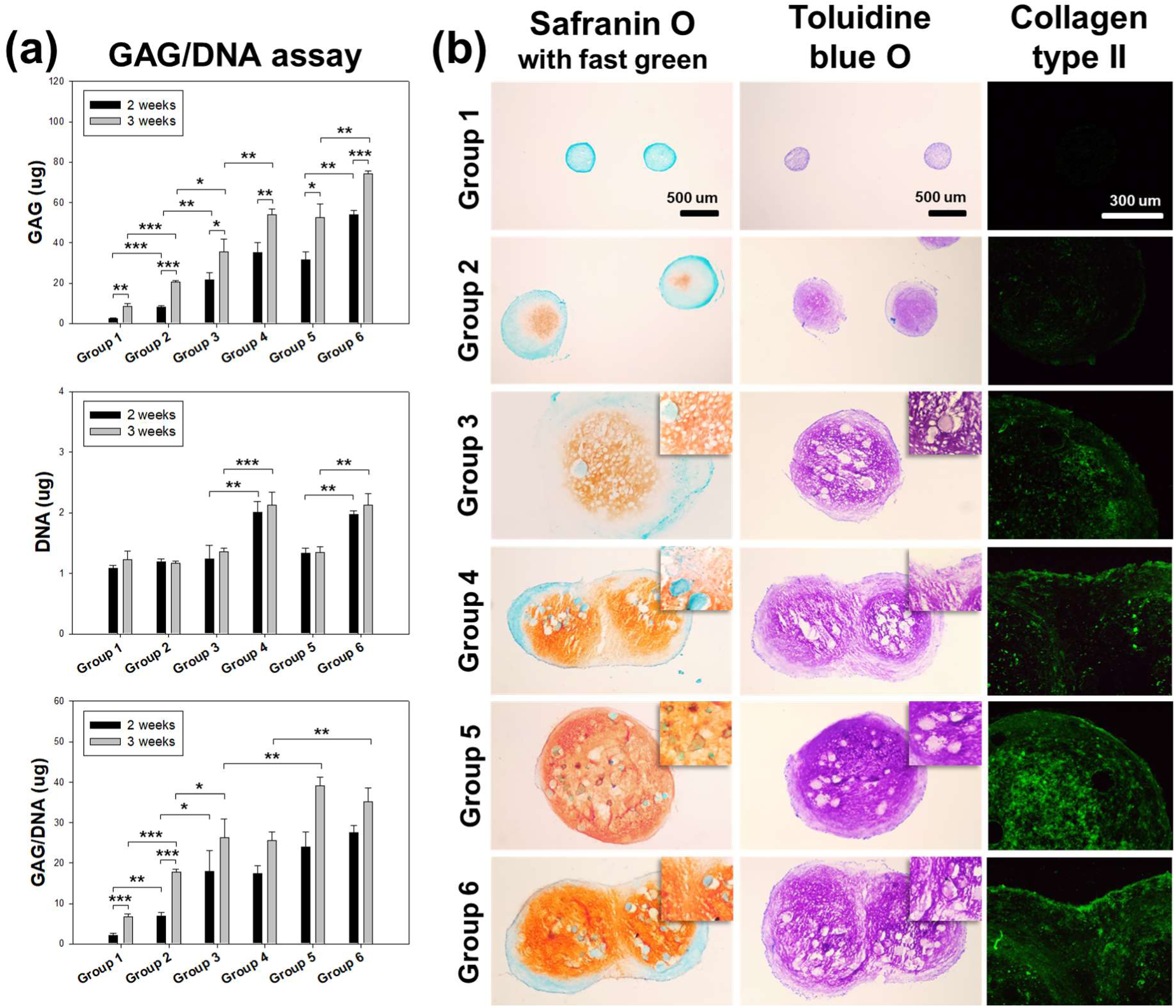
*In vitro* chondrogenesis of hMSC and hMSC/GMs core / 4-OMA shell microgels in different media conditions. (a) Quantification of GAG, DNA, and GAG/DNA content for 2 and 3 weeks of culture and (b) photomicrographs of tissue sections stained with safranin O with fast green, toluidine blue O, and immunofluorescence against collagen type II after 3 weeks of culture. Small insets show GMs embedded within condensations and the fusion area between paired microgels. The GAG/DNA contents and the staining intensity in TGF-β1-releasing GMs encapsulated microgels were higher than those with exogenous TGF-β1 supplied in the media. Both aggregate fusion and chondrogenic differentiation were excellent with TGF-β1 supplied via GM within the core of the microgels. In the statistical analysis of GAG and GAG/DNA levels at 3 weeks, groups 5 and 6 were significantly higher than groups 3 and 4, respectively (*p\** < 0.05, *p\*\** < 0.01, *p\*\*\** < 0.001).

Histologic sections were stained with Safranin O/fast green, toluidine blue O, or collagen type II to identify the presence of GAG and the distribution of collagen within the 3D condnesations after 3 weeks (Figure 4b). Groups with exogenously supplemented TGF-β1 or TGF-β1-releasing-GMs (groups 3-6) exhibited more intense staining than groups 1 and 2. The embedded empty GMs and TGF-β1-releasing GMs were relatively evenly distributed throughout the cellular condensations (small inset). In the double aggregates (groups 4 and 6), the cellular condensations were well fused with strong chondrogenic differentiation. The most intense staining was demonstrated in groups 5 and 6. These histological results corroborated the GAG quantification results.

### 2.4 Degradation of core/shell microgels with encapsulated cells *in vivo*

hMSC-laden core/shell microgels were subcutaneously implanted into mice without prior *in vitro* culture, and then harvested at 3 and 7 days post-surgery (Figure 5). In the unmodified alginate group, the original round microgel geometry was observed at harvest (blue arrows). The residual unmodified alginate was visualized as an acellular area (black stars) in the alcian blue with nuclear fast red and hematoxylin/eosin (H&E) staining. Encapsulated hMSCs formed condensations within the core area (yellow stars), while the residual acellular unmodified alginate impaired microgel fusion with host tissue. In the 4-OMA core/shell microgel group, the alginate shell layer was completely degraded at both harvest time points, and the implanted hMSCs formed condensations and integrated with themselves and host tissues (red arrows). In the H&E and alcian blue staining, implanted hMSCs (blue stars) seamlessly integrated with host adipose tissues (red stars).

**Figure 5.**
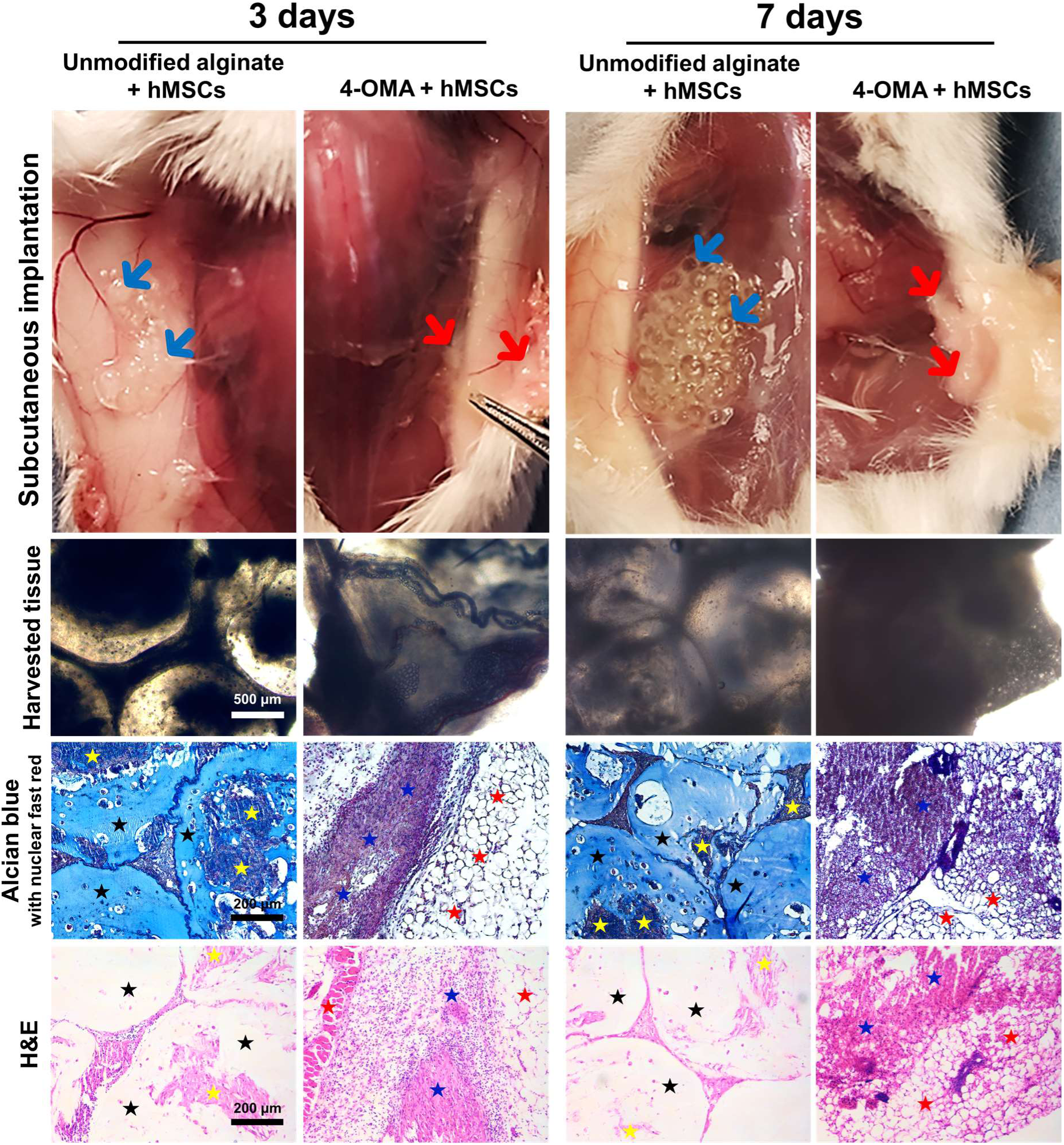
Subcutaneous *in vivo* characterization of tissue fusion between implanted core/shell microgels and host tissue 3 and 7 days post-surgery. The hMSC encapsulated core / unmodified alginate or 4-OMA shell microgels were implanted in subcutaneous pouches on the backs of CB-17 SCID mice without prior *in vitro* culture. The presence and absence of residual alginate, core cell condensation formation, and tissue fusion were visualized by macroscopic and microscopic images. The alcian blue with nuclear fast red and H&E staining showed the persistence of residual unmodified alginate, which interfered with condensation fusion and integration with surrounding tissue, while there was not residual alginate in the 4-OMA shell group, and hMSCs in the core condensed, fused with each other, and integrated with surrounding host tissue (blue arrows: residual alginate; red arrows: hMSC condensations integrated with each other and host tissue; yellow stars: implanted hMSCs in unmodified alginate group; blue stars: implanted hMSCs in 4-OMA microgel group; red stars: host adipose tissue including blood vessels).

### 2.5 *In vivo* chondrogenesis of immediately implanted core/shell microgels

hMSC-laden core/shell microgels were subcutaneously implanted into mice, and then harvested at 1 and 3 weeks post-implantation. Detailed descriptions of the *in vivo* experimental groups for driving chondrogenesis are described in Table. S3. At 1 week post-surgery, group 1 (hMSCs and TGF-β1-releasing GMs core / unmodified alginate shell microgels) did result in condensation of the hMSCs in the microgels (Figure 6a). The residual unmodified alginate, stained a strong blue color (black stars) by the dye, alcian blue, interfered with tissue fusion. In contrast, groups 2 (hMSCs and empty GMs core / 4-OMA shell microgels) and 3 (hMSCs and TGF-β1-releasing GMs core / 4-OMA shell microgels) did exhibit hMSC condensation formation as well as tissue fusion. The implanted hMSCs exhibited strong integration with host muscle and adipose tissues (red stars) as visualized via H&E and alcian blue staining. This result was further confirmed by human nuclear antigen (HNA) immunohistochemistry staining (Figure 6b). The hMSCs were positively stained by HNA (red arrow) in the transplanted constructs, but the host tissue control was negative for HNA staining.

**Figure 6.**
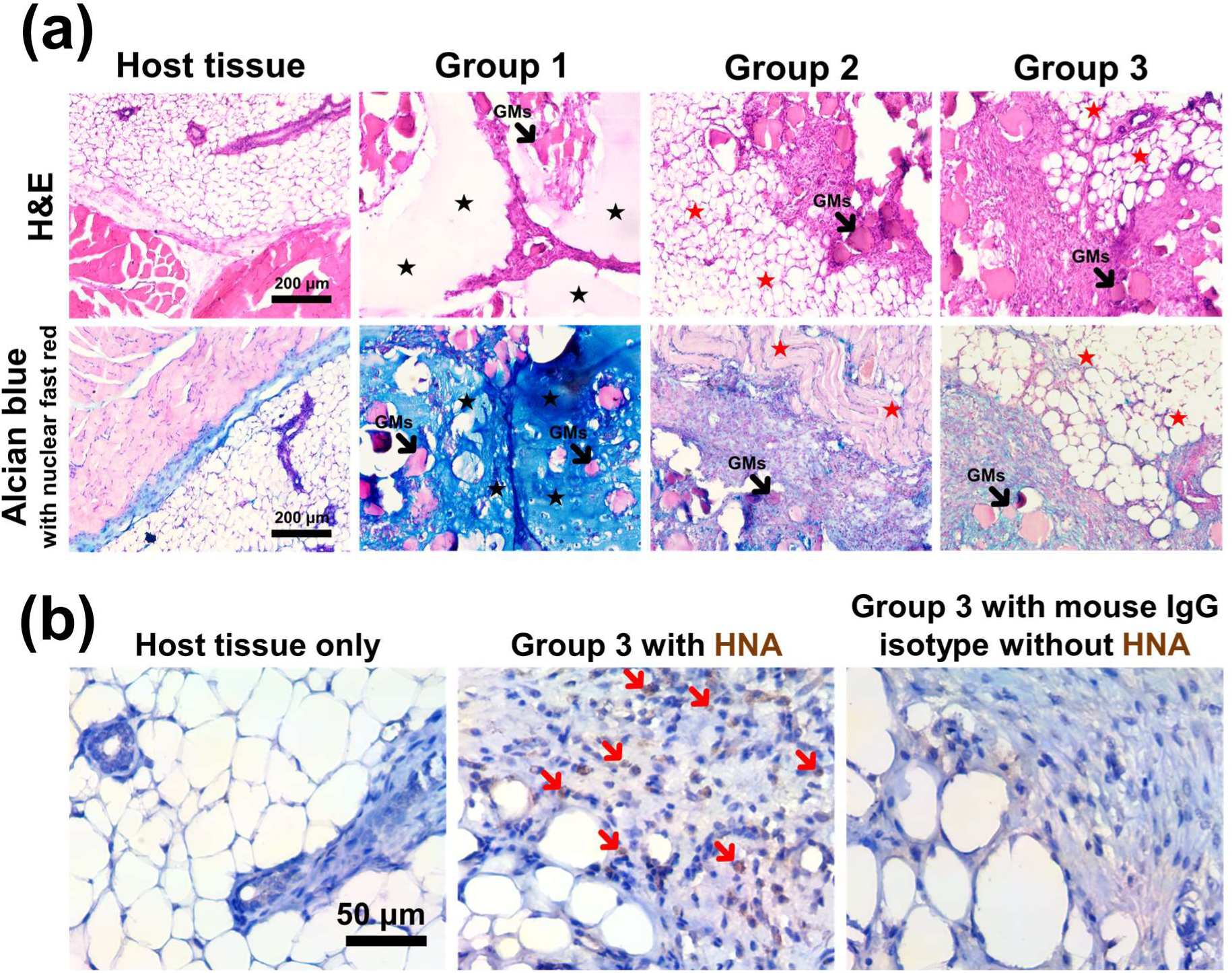
Subcutaneous *in vivo* characterization of tissue fusion between implanted hMSCs & GM core / shell microgels and host at 1 week post-surgery. (a) Tissue fusion was visualized by H&E, and alcian blue with nuclear fast red staining. In all histological staining, the GMs were distributed relatively uniformly within the formed core hMSC condensations. The hydrogel component remained in group 1, whereas tissue integration between implanted hMSCs condensations and host tissue was demonstrated in groups 2 and 3. (b) Immunohistochemistry staining for HNA was performed to visualize hMSCs within the mouse tissue. The hMSCs nuclei stained positively (red arrows) and their extracellular matrix was well integrated with mice host tissue. The host tissue with primary-HNA antibody and group 3 with mouse IgG isotype without primary-HNA antibody were not positively stained (black stars: residual alginate; red stars: host muscle and adipose tissues; black arrows: GMs; red arrows: hMSCs stained by HNA)

At 3 weeks post-implantation, unmodified alginate shell-based microgels retained their original round shape while the 4-OMA shell-based microgels completely degraded (Figure S8). Group 1 displayed a cartilage-like matrix with differentiated chondrocytes within lacunae (blue arrow) surrounded by dense cartilage matrix (Figure 7a). The acellular residual alginate (black stars) impaired tissue fusion between the generated cartilage matrix and surrounding host tissue. In group 2, core cellular condensation tissues with empty GMs fully integrated without residual alginate. However, this group only displayed formation of connective tissue, and not cartilage matrix. In group 3, abundant cartilage-like matrix formed with chondrocytes within lacunae (blue arrow) that integrated with host tissue. Similar staining results were observed in other replicate animals (Figure S9-12). GAG and GAG per DNA levels of group 1 and 3 were significantly higher than those of group 2 (Figure. 7b), while there was no significant difference in GAG production between groups 1 and 3.

**Figure 7.**
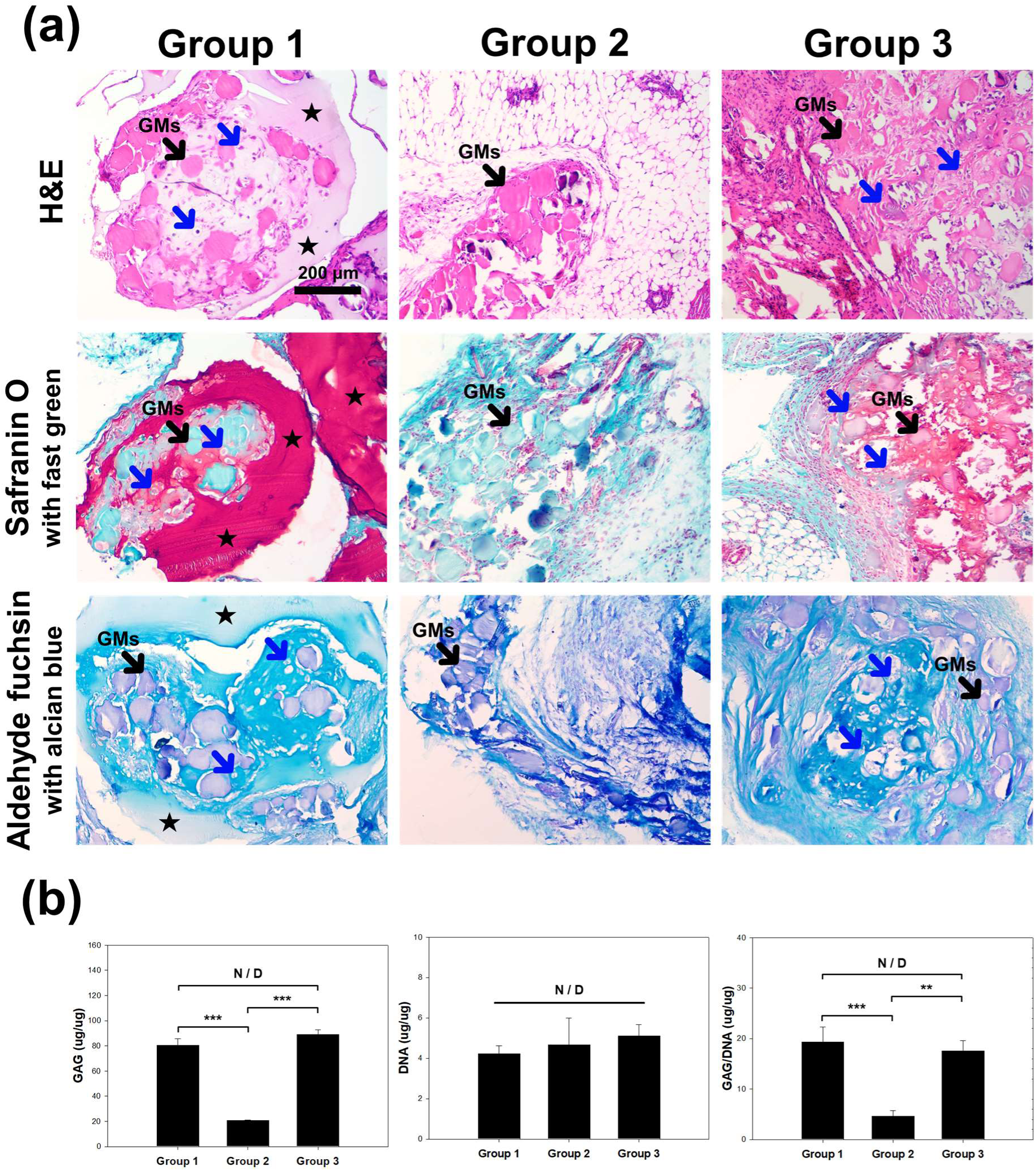
Characterization of *in vivo* subcutaneous chondrogenic differentiation of hMSC/GMs encapsulated core / 4-OMA shell microgels at 3 weeks post-surgery. (a) Photomicrographs of histological sections after 3 weeks. In group 1, unmodified alginate remained and impaired tissue integration with host tissue but cartilage tissue was generated within the alginate shell. The residual alginate is seen as an acellular area. In group 2, the 4-OMA shell degraded and tissue fusion was evident but there was no chondrocytes or cartilage matrix readily visible. In the group 3, the alginate shell also degraded and robust cell condensation-based cartilage-like tissue formed. Group 2 and 3, but not group 1, had strong integration with host tissue. (b) GAG, DNA, and GAG/DNA content after 3 weeks. Groups 1 and 3 exhibited significantly higher GAG and GAG/DNA content compared to Group 2. There was no significant difference between groups 1 and 3 in GAG and GAG/DNA content. There was no significant difference in DNA content between groups. (black stars: residual alginate; black arrows: residual GMs; blue arrows: chondrocyte in lacunae;).

### 2.6 Further *in vivo* characterization of engineered cartilage matrix along with surrounding host tissue

Differentiated chondrocytes derived from hMSCs were visualized by HNA staining (Figure S10b). In group 1, differentiated chondrocytes within cartilage matrix were positively stained (blue arrows in “i”). However, the unmodified alginate still remained and hindered tissue fusion. In group 3, cells of host murine tissue did not stain for HNA (purple arrows in “ii”), but the differentiated chondrocytes in adjacent cartilage-like tissue were positively stained (blue arrows in “ii”). In the central area of abundant neocartilage, the chondrocytes were embedded within a cartilage-like matrix (blue arrows in “iii”).

### 2.7 Formation of assembled structures with defined architectures by core/shell microgel manual assembly and photocrosslinking

This system may also be used to manually assembled larger defined complex architectures for subsequent fusing of cell condensations into structures with specific geometries. After manually connecting individual core / 4-OMA shell microgels on a petri dish, triangle and rhombus structures were assembled by photocrosslinking of the core/shell microgels (Figure 8a). Because the thin layer of 4-OMA has methacrylate groups, it enabled photocrosslinking between the microgels to stabilize the assemblies (Figure 8b).^[21]^ The 4-OMA shell layer, when irradiated with low level UV light, formed physical linkages between the microgels. The high-density hMSCs were observed within the 4-OMA shell layers. These high-density hMSCs-laden core/shell microgels could be directly assembled while maintaining their original core/shell structure. It was demonstrated that photocrosslinked robust constructs maintained their original shape under strong pipetting in the PBS (Figure S Movie 3).

**Figure 8.**
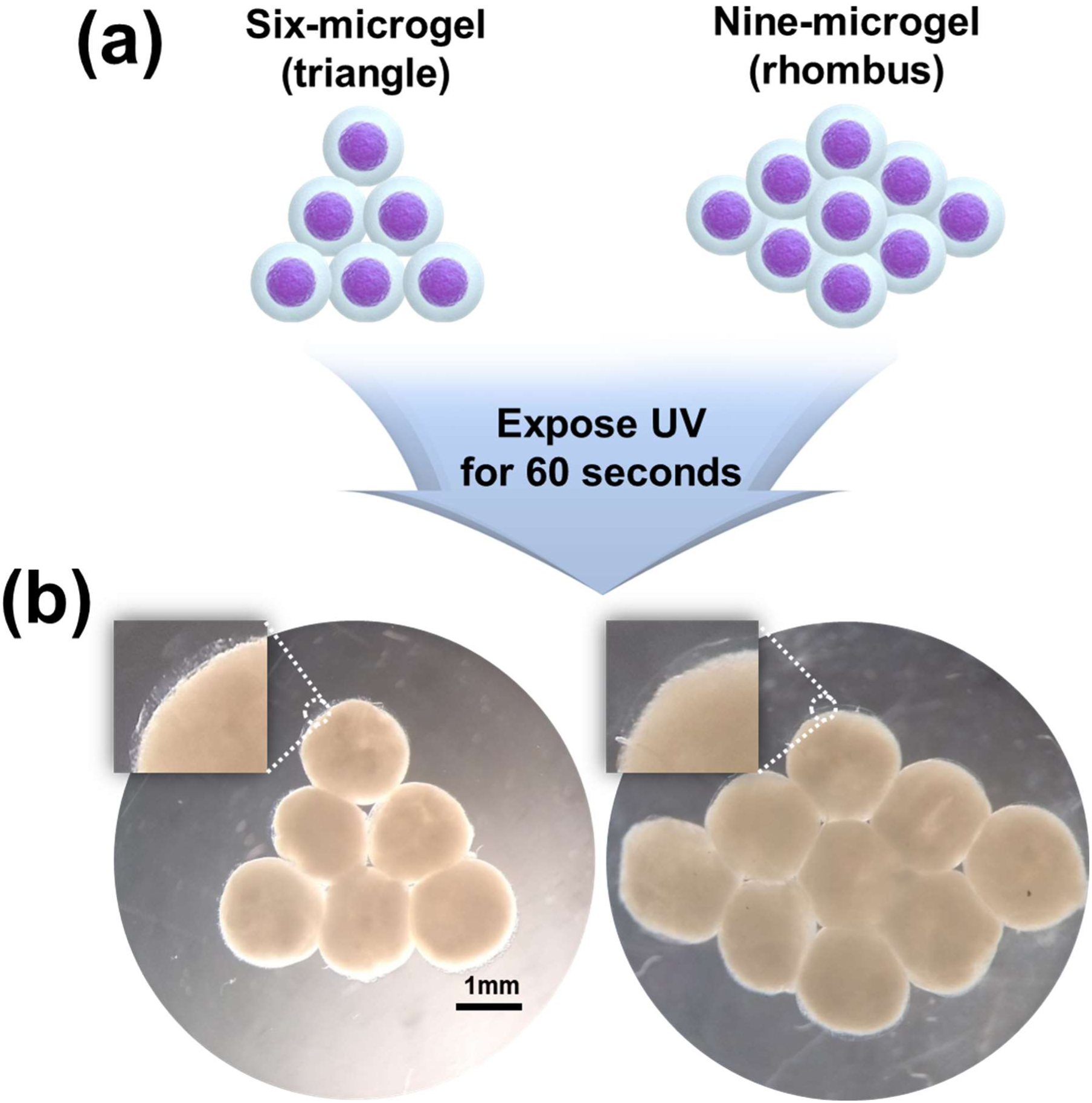
Formation of assembled structures with defined architectures by core/shell microgel manual assembly and photocrosslinking. (a) NIH3T3 core / 4-OMA shell microgels were manually organized as triangle and rhombus shapes on a petri-dish, and then photoinitiator containing medium was added to the constructs followed by UV exposure. (b) Then, the constructs were visualized via microscope in the petri dish. Both constructs robustly maintained their original architecture. Thin 4-OMA shell layers served as physical connections between microgels (b, small inset).

## 3. Discussion

Clinically, delayed treatment of damaged cartilage tissue may result in further degeneration and predictably poor outcomes.^[22]^ Indeed, cartilage injuries and osteoarthritis can have serious negative effects such as subchondral bone resorption,^[23]^ the development of bone marrow edema,^[24]^ and chronic pain.^[25]^ While in some circumstances rapid treatment for damaged cartilage tissue may be valuable to prevent ensuing serious osteochondral complications,^[26]^ unfortunately, there is no current method of reliably forming new articular cartilage to fill these defects with demonstrated long-term clinical benefit.

Cell condensation-based strategies are promising for cartilage tissue engineering since they allow for abundant cell-cell contacts mimetic of cartilage formation during skeletal development.^[27]^ Microwell systems and hanging drop techniques have been commonly used for the generation of cell condensation-based cartilage tissue,^[28]^ but prolonged *in vitro* culture period is required due to the need to provide geometric confinement for condensation formation (e.g., base/walls of microwell, surface tension of liquid drop) and chondroinductive factors in cell culture media to induce chondrogenic differentiation. Another strategy involving stirring of a cell suspension in a spinner flask^[29]^ results in non-uniform 3D cell condensations and also requires a prolonged period of *in vitro* culture time with a substantial amount of expensive medium. All of these methods cannot be used as an immediate implantation therapy due to the *in vitro* culture period required to achieve cell condensations and induce their maturation into neocartilagenous tissues.

Our core/shell technology utilized in this study enables the engineering of cell condensations within the core of the microgels, promoting cartilage formation. This approach involves encapsulating cells within a core material while providing a protective shell layer to prevent cell dispersion after transplantation but prior to condensation formation. Other research groups have explored engineering constructs to generate chondrogenic cell condensations. Loo et al. developed hollow compartmentalized hydrogel encapsulating cellular microspheroids via in air microfluidic system for ultimate generation of biomaterial-free chondrogenic cellular tissues.^[30]^ They successfully produced high-throughput spheroids in hydrogels, but the surrounding alginate required manual removal enzymatically using an alginate lyase washing step after *in vitro* culture. Importantly, it was also necessary for the cellular spheroids within the hydrogels to be cultured *in vitro* in chondrogenic medium for 21 days for condensation formation and chondrogenesis. In another study, Caprio et al. combined cell spheroids and similarly-sized microgel particles to form granular composites.^[31]^ Spheroid fusion and cartilage tissue formation occurred, but required *in vitro* culture in the presence of chondrogenic factors.^[13, 18b]^ While these studies successfully achieved 3D chondrogenic condensations they required *in vitro* culture, manual removal of used hydrogels used and/or the addition of exogenous chondroinductive signals for differentiation. For these reasons, it would not be possible to implant the constructs immediately in the body and achieve cell condensation and cartilage tissue formation AND fusion and integration of the resulting microtissues with each other and surrounding tissue.

In contrast, our core/shell microgels offer multiple critical advancements all within the same system. First, the core of the microgels provides the necessary space for cellular self-assembly into condensations and subsequent formation of cartilage-like tissue within the engineered construct (Figure 4). The shell layer of the microgels contains the core cell contents long enough for condensation formation, but then hydrolytically degrades rapidly over 2-3 days, facilitating *in vivo* fusion and integration of the condensations with each other and surrounding host tissue (Figure 5). This shell degradation addresses a key limitation of non-degrading shells in other systems.^[32]^ Constructs may be immediately implanted, and cells can form condensations that can fuse rapidly since the shell degrades quickly. Moreover, this system eliminates the need for exogenous chondroinductive signals, further enabling immediate implantation. In contrast to other approaches, inclusion of chondroinductive TGF-β1-releasing GMs with the cells within the cores of the microgels drives local chondrogenic differentiation, leading to the directed formation of cartilage-like tissue *in vivo* (Figure 7, S9-12). This eliminates the requirement for extensive *in vitro* culture before implantation and reduces the amount of expensive growth factors needed.^[10]^ The key findings from our study demonstrate the successful condensation of implanted cells, the rapid degradation of the shell layer, and the formation and integration of functional cartilage-like tissue without prior *in vitro* culture (Figure 7, S9-12). This biomaterial degradation is crucial for engineered tissue integration, maturation and remodeling, and for the long-term success of engineered cartilage.^[33]^

Furthermore, our ability to create complex architectures using the core/shell microgels opens opportunities for developing patient-specific cartilage replacements with tailored geometries (Figure 8). Based on the present system’s novel ability to coordinate biomaterial removal and tissue formation, it may open new avenues for cartilage tissue reconstruction.

## 4. Conclusion

In summary, this work presents an innovative high-density hMSCs core / hydrogel shell delivery system for use in immediate implantation therapy. Through chemical modification of the alginate, quickly and fully degradable OMA was obtained, which is cyto- and biocompatible. Upon fabrication, the core/shell constructs could be cultured *in vitro* or implanted immediately *in vivo*, with the hydrogel shell persisting long enough to allow for the formation of 3D hMSC cell condensations. When chondroinductive TGF-β1-releasing GMs were incorporated into the core, the growth factor drove hMSC chondrogenic differentiation and the formation of cartilage-like tissue *in vitro* and *in vivo*. Importantly, the hydrogel shell was designed to degrade rapidly by hydrolysis, enabling spontaneous fusion of adjacent cellular constructs *in vitro* and additionally excellent integration with host tissue and effective chondrogenesis without any prior *in vitro* culture *in vivo*. It may be possible to use this system to form gradient tissue constructs mimicking, for example, bone-to-cartilage tissue transition via UV irradiation-mediated core-shell particle fusion to reconstruct damaged osteochondral defects.^[34]^ This core/shell microgel system may have great potential for rapid therapeutic stem cell delivery for stem cell condensation-based cartilage tissue engineering and for other tissue engineering and regenerative medicine applications.

## 5. Materials and methods

A detailed description of the materials, analysis, and experimental methods are presented in the Supplement Information.

## Acknowledgements

The authors wish to express thanks to Mrs. Dasom Seo for preparation of schematic illustrations. The authors gratefully acknowledge funding support from the Department of Veterans Affairs, Veterans Health Administration, Office of Research and Development, Rehabilitation Research and Development Service under award number RX004288 and the National Institutes of Health’s National Institute of Arthritis and Musculoskeletal and Skin Diseases under award number R01AR081448. The contents of this publication are solely the responsibility of the authors and do not necessarily represent the official views of the Department of Veterans Affairs or the National Institutes of Health.

---

This study reports an innovative high-density hMSCs core / hydrogel shell delivery system for use as an immediate implantation therapy. To do this, quickly and fully degradable oxidized and methacrylated alginate (OMA) was obtained. Upon fabrication, the core/shell constructs could be implanted immediately in vivo, with the hydrogel shell persisting long enough to allow for the formation of 3D hMSC cell condensations and chondrogenic differentiation, and the formation of cartilage-like tissue *in vitro* and *in vivo*. This core/shell microgel system may have great potential for rapid therapeutic stem cell delivery for stem cell condensation-based cartilage tissue engineering and for other tissue engineering and regenerative medicine applications.

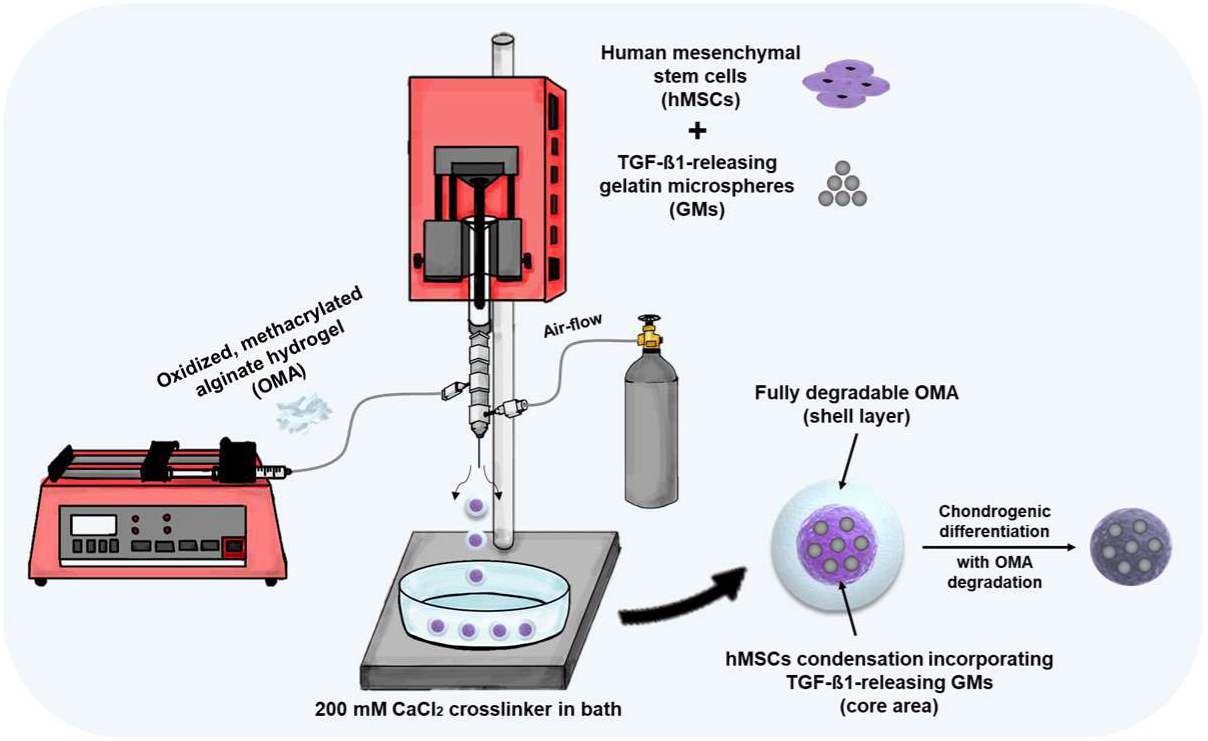
ToC figure.

