## Supporting information for "*In situ* cell condensation-based cartilage tissue engineering via immediately implantable high-density stem cell core and rapidly degradable shell microgels"

### 2. Materials and methods

2.1. Materials for preparation of oxidized and methacrylated alginate (OMA)

Sodium alginate (Protanal® LF120M, 157 mPas) was generously provided as a gift from NovaMatrix/FMC Biopolymer (Sandvika, Norway). Sodium periodate ($NaIO_4$, Cat. No. 311448, ACS reagent, ≥99.8%) and MES hydrate (Cat. No. M8250, ≥99.5% (titration)) were purchased from Sigma-Aldrich (St. Louis, MO). Sodium chloride (NaCl, Cat. No. S271-1), calcium chloride dihydrate ($CaCl_2$, Cat. No. C79-500), and Tween®20 (Cat. No. BP337-500) were purchased from Fisher Scientific (Pittsburgh, PA). 2-Aminoethyl methacrylamide hydrochloride (AEMA, Cat. No. 21002-10) was purchased from Polysciences Inc. (Warrington, PA). N-hydroxysuccinimide (NHS, 98%) was purchased from Acros organics (Fair Lawn, NJ). 1-Ethyl-3-(3-dimethylaminopropyl) carbodiimide hydrochloride was purchased from ProteoChem (Denver, CO). Spectra/Por® biotech cellulose ester dialysis membranes (3,500 MWCO) were purchased from Spectrum Laboratories, Inc. (Rancho Dominguez, CA). Activated charcoal (Cat. No. 05-690b, 50-200 mesh) was purchased from Fisher Scientific (Pittsburgh, PA).

2.2. Preparation of OMA

The OMA was synthesized by oxidation and methacrylation processes as described in our previous work with some modifications.[1] Briefly, sodium alginate (10 g) was dissolved in ultrapure deionized water ($diH_2O$, 900 ml) overnight. Sodium periodate (0.216 and 0.432 g) was dissolved in 100 ml $diH_2O$ separately, and then added into respective alginate solutions to attain different degrees of theoretical alginate oxidation (2 and 4 %, respectively) under vigorous stirring in the dark (covered by aluminium foil) at room temperature for 24 h. For the methacrylation of oxidized alginate, MES hydrate (19.52 g) and NaCl (17.53 g) were directly added to an oxidized alginate solutions (1L) and then the pH was adjusted to 6.5. After that,

NHS (2.65 g) and EDC (8.75 g) were added to the mixtures under stirring to activate carboxylic acid groups of the alginate. After 5 min, AEMA (3.376 g, theoretical 40 % methacrylation of oxidized alginate) was added into oxidized alginate solutions very slowly and the reactions were maintained in the dark with stirring at room temperature for 24 h. The reacted mixtures were precipitated into excess acetone, dried in a fume hood, and rehydrated to 1 % w/v solutions in $diH_2O$ for further purification. The OMAs were purified by dialysis membrane in $diH_2O$ for 3 days. After that, they were treated with activated charcoal (0.5 mg/100 ml) for 30 min and then filtered using bacteria excluding 0.22 μm pore size Steritop® filters (Millipore GP Express, Billerica, MA). Finally, the products were lyophilized at least 10 days, and then stored at 4 °C until usage. The theoretical 2 and 4 % oxidized, 40 % methacrylated alginate were defined as 2-OMA and 4-OMA, respectively. To verify the actual oxidation degree via $^1$H-NMR analysis, part of oxidized alginate solution was dialyzed and lyophilized prior to the methacrylation procedure.

2.3. Preparation of $CaCl_2$ cross-linker

For the crosslinking of core/shell hydrogel particles, cross-linker was prepared using $CaCl_2$. Briefly, 25 mM MES hydrate (9.762 g) was completely dissolved in ultrapure deionized water ($diH_2O$, 2000 ml). After that, 150 mM $CaCl_2$ (44.103 g) was added into the mixture and the pH was adjusted to 7.2. The mixture was filtered using bacteria excluding 0.22 μm pore size Steritop® filters (Millipore GP Express, MA). Finally, 0.05 wt. % Tween® 20 was added into the mixture. The manufactured cross-linker was kept at room temperature.

2.4. Preparation of TGF-β1-releasing gelatin microparticles (GMs)

TGF-β1-releasing GMs were prepared as previously described with slight modification.[2] Briefly, 5 g of gelatin from porcine skin (gel strength 300, Type A, Cat. No.

G2500) was dissolved in 45 ml of di$H_2O$ under stirring at 60 °C. The gelatin solution was slowly (2 drops / sec) added dropwise into 250 ml of preheated olive oil at 45 °C under stirring at 500 RPM for 10 min. The solution temperature was lowered to 15 °C with constant stirring to facilitate gelation by putting the suspension in a 4 °C fridge. After 30 min, 100 ml chilled acetone (4 °C) was added to the mixture under stirring. After 1 h, an additional 100 ml of acetone was added to the mixture and stirred for 5 min at 1000 RPM. The produced GMs were then harvested by kimwipe filtration with vacuum followed by washing with acetone three-times to remove residual oil. Finally, the GMs were air dried in a fume hood.

Dried GMs were cross-linked at room temperature in an aqueous solution of 1 wt. % genipin (Cat. No. 078-03021, Wako Pure Chemical Industries, Osaka, Japan). Briefly, 1g of GMs was added into 100 ml of 1 wt. % genipin solution under stirring at room temperature for 4 h. Cross-linked GMs were harvested by kimwipe filtration with vacuum, washed 3 times with di$H_2O$, and lyophilized. The TGF-β1 loaded 1 mg of GMs were prepared by immersing cross-linked, ultraviolet (UV)-sterilized GMs in 400 ng/5 ul of TGF-β1 (Cat. No. 100-21, PeproTech, Rocky Hill, NJ) solution in phosphate-buffered saline (PBS) for 2 h in a 37 °C water bath immediately before use. The empty GMs were hydrated similarly using PBS only.

2.5 Preparation of core/shell microgels via triaxial nozzle system

To fabricate core/shell microgels, we designed a triaxial nozzle system (Figure S1). Injections using two syringe pumps (Model No. NE-1000X, New Era Pump Systems Inc., Farmingdale, NY) and airflow were all carried out simultaneously into a $CaCl_2$ bath (Figure S Movie 1). A prebuilt commercial triaxial nozzle (Cat. No. 100-10-TRIAXIAL-241814) was purchased from Ramé-hart Inc. (Succasunna, NJ).

To set up the core/shell microgel generation system, a 1 ml gastight syringe (Cat. No. 1001 TLL, Hamilton Co, Reno, NV) was filled with PBS, NIH3T3s, hMSCs, hMSCs/empty

GMs, or hMSCs/TGF-β1-releasing GMs, depending on the experimental group, and then the pre-filled 1ml syringe was connected to the core part of needle.

For the middle part of needle that is used to form the shell, 2 wt. % of unmodified alginate, 2-OMA and 4-OMA were separately thoroughly dissolved in PBS at room temperature, and then warmed in a 37 °C water bath before use. After preparation of the macromer solutions, a 3 ml BD plastic luer-lock™ disposable syringe (Cat. No. 14-823-435, Fisher Scientific, Pittsburgh, PA) was filled with either unmodified alginate, 2-OMA or 4-OMA solution. The middle part of the needle was connected to the macromer solution pre-filled 3ml syringe through PVC tubing (180 metric clear, Nalgene®, 8001-0204, Rochester, NY) and male/female syringe tip connectors ($^3/_{32}$ inch, McMaster-Carr, Atlanta, GA).

The outer part of the needle was connected to air-flow using a gas flow meter (Model: PMR1-010973, Cole-Parmer Instruments Inc., Vernon Hills, IL) through PVC tubing and male/female syringe tip connectors. Finally, Finally, the two syringes were attached to syringe pumps.

The injection speed of the syringes was fixed (core: 99 ul/min, shell: 55 ul/min), but airflow was modulated over a range by varying the air pressure from 15.5 to 17 psig to obtain different sizes of microgels (Figure 2c and d) (n=4).

### 2.6 Degradation test of core/shell microgels with/without cells

For the degradation test of OMA, we used 1,9-dimethylmethylene blue (DMMB), a cationic dye that can bind with negatively charged polysaccharides such as alginate molecules.[3] In this assay for quantifying alginate, a DMMB dye solution at pH 7 was used because it binds to alginate and other GAGs like those in cartilage at pH above 3.[3a] DMMB powder (10.5 mg) was dissolved in 2.5 ml absolute ethanol, and then 1 g of sodium formate (ACS reagent, ≥99.0%, Sigma-Aldrich) was added. The volume was brought up to 450 ml by adding ultrapure deionized water, and then the pH was adjusted to 7 using a formic acid solution (BioUltra, 1.0 M in $H_2O$, Sigma-Aldrich). Finally, ultrapure deionized water was added to bring the volume to 500 ml. The DMMB solution was wrapped with aluminum foil and stored at 4 °C until use.

To verify the degradation kinetics without cells, we generated core/shell microgels using 2- and 4-OMA. The core part was filled with PBS only. The generated core/shell microgels were randomly picked and incubated in 96-well clear round bottom ultra-low attachment microplates (Cat. No. 7007, Corning®, NY) with general growth medium for 48 h at 37 °C and 5 % $CO_2$. The supernatant medium was carefully harvested at 3, 12, 24, and 48 h (n=4, per time point). The content of degraded OMA in the supernatant medium was measured by the DMMB assay. To do this, 100 µl of supernatant medium and 100 µl of DMMB solution were mixed vigorously in clear 96 well plates. The well plates were covered by aluminum foil to prevent light entry and then incubated at room temperature for 30 min. Absorbance intensities of the samples were recorded at 520 nm using a microplate reader (SpectraMax® iD3, Molecular Devices).[3b, 4]

To measure the degradation of hydrogels with cells, high-density cells encapsulated core/shell 2- and 4-OMA microgels were generated. The detailed *in vitro* cell culture process is detailed in section 2.8. The core part was filled with mouse fibroblast NIH3T3 cells. The NIH3T3 encapsulated core/shell microgels were randomly picked and incubated in 96-well clear round bottom ultra-low attachment microplates (Cat. No. 7007, Corning®, NY) with cell culture growth medium for 72 h at 37 °C and 5 % $CO_2$. The supernatant media was carefully harvested at 3, 12, 24, 48, and 72 h (n=5, per time point). After supernatant were harvested, to measure the remaining OMA within the cell aggregates, they were placed in Eppendorf Tubes® (EP-Tubes®) with cell culture growth medium and then pulverized with a homogenizer (TH115-PCR-D; Omni International Inc., Marietta, GA). Finally, the suspensions were centrifuged at 300 x g for 5 min to get supernatants only without cell debris. The ensuing remaining alginate measurement process was the same as that used on core/shell microgels without cells.

2.7 $^1$H-NMR analysis

20 mg of unmodified alginate, oxidized alginate, or OMA were dissolved in 1ml of $D_2O$ (Cat. No. 450510, Sigma-Aldrich, St. Louis, MO). $^1$H-NMR spectra were acquired with an AVANCE III HD 400 MHz spectrometer (Bruker BioSpin Corp., Ettlingen, Germany) operating at 25 °C.

2.8. *In vitro* analysis

2.8.1. Preparation and expansion of NIH3T3 and hMSCs

In this study for biological assessments, mouse fibroblast cells (NIH3T3) were used for system modelling *in vitro* and human bone marrow-derived mesenchymal stem cells (hMSCs) were used for chondrogenesis experiments with cartilage tissue formation *in vitro* and integration additionally *in vivo*. The hMSCs were isolated as previously described.[2, 5] To use either cell type for experiments, NIH3T3 or hMSCs were expanded in cell culture growth medium consisting of low glucose-Dulbecco's modified Eagle medium (LG-DMEM, Sigma-Aldrich) with 10 % fetal bovine serum (FBS, Sigma-Aldrich), and 1 % penicillin-streptomycin (P/S, Gibco®, Invitrogen, Grand Island, NY) in an incubator at 37 °C and 5 % $CO_2$. 10 ng/ml of recombinant human fibroblast growth factor-2 (rhFGF-2, Cat. 233-FB, R&D Systems, Minneapolis, MN) was also used in the hMSC expansion medium. Expanded cells were harvested with 0.05 % trypsin/EDTA (Thermo Fisher Scientific, Waltham, MA) and concentrated by centrifugation at 300 x g for 5 min. After aspiration of the supernatant, cell pellets were loaded in a 1 ml gastight syringe to use as a cell-only core encapsulated in a hydrogel shell layer. For a comparison with the core/shell microgel system, 3D cell spheroids without a hydrogel shell were prepared by a standard centrifugation method using a 96-well V-bottom polypropylene plate (Cat. 651201, Greiner Bio-One, Frickehausen, Germany) with a polypropylene microplate lid (Cat. 290-8020-03L, Caplugs Evergreen, Buffalo, NY).[6] After 7 days of culture, the diameter of the 3D spheroids was measured (n=4) based on scales from microscope images. The x-axis and y-axis diameters of 3D spheroids were measured and averaged using Microsoft Windows 10 Paint. Diameters of 3D aggregates formed from 4-OMA core/shell microgels were compared to those formed from the centrifugation in V-bottom plate method.

2.8.2. Characterization of NIH3T3 encapsulated core/shell microgel and evaluation of fusion

For further *in vitro* analysis, NIH3T3 encapsulated unmodified alginate and 2- and 4-OMA core/shell microgels were formed, and then individual microgels were cultured in 96-well clear round bottom ultra-low attachment microplates (Cat. No. 7007, Corning®, NY) with cell culture growth medium at 37 °C and 5 % $CO_2$ for 21 days. To evaluate core cell aggregate fusion, two core/shell hydrogel particles were cultivated together in the same 96-well space as soon as they were formed. The morphology of individual single core/shell microgels and tissue fusion of double core/shell microgels were evaluated using inverted phase-contrast microscopy (TMS-F, Nikon Corporation, Tokyo, Japan) for 21 days and 3 days, respectively.

To measure DNA content, NIH3T3 encapsulated 2- and 4-OMA core/shell microgels were cultured for 21 days. During culture, cell aggregates were harvested and stored in a -20 °C freezer at a predetermined time points up to 21 days. After harvest, all samples were digested in 1 ml of papain buffer solution (25 μg/mL papain, 2 mM L-cysteine, 50 mM sodium phosphate, 2 mM ethylenediaminetetraacetic acid) at 65 °C overnight. Finally, DNA content was measured at 0, 1, 3, 7, 14, and 21 days post-culture via Quant-iT™ PicoGreen® (Invitrogen,

Carlsbad, CA) assay according to the manufacturer's instruction (n=4).[1, 5a] For live/dead staining, NIH3T3 encapsulated 4-OMA core/shell microgels were cultured for 7 day. Before staining, diluted live/dead solution was prepared. Briefly, fluorescein diacetate (FDA, 1.5 mg/ml in dimethyl sulfoxide; stains live cells green) and propidium iodide (PI, 1 mg/ml in PBS; stains dead cells red) solutions were mixed with PBS at a diluted ratio (1:500). After washing cell aggregates three time with PBS, 100 µl of staining solution was added into each 96-well and incubated for 10 min at room temperature, and then stained cell aggregates were visualized using a fluorescence microscope (ImageXpress® Pico, Molecular Devices, San Jose, CA). Portions of the sample views were stitched together into a single mosaic images due to size limitations (n=4, but only 1 shown as all appeared similarly stained).

To fuse the shells of core-shell building-blocks together into complex architectures (Figure 8), NIH3T3 encapsulated 4-OMA core/shell microgels manually organized into triangle and rhombus geometries were immersed in high glucose-Dulbecco's modified Eagle medium (HG-DMEM, Sigma-Aldrich) containing 0.05 wt. % 2-Hydroxy-4'-(2-hydroxyethoxy)-2-methylpropiophenone photoinitiator (Cat. No. 410896, Sigma-Aldrich) for 15 s. After that, organized constructs were exposed to UV light (320–500 nm, EXFO OmniCure S1000-1B, Lumen Dynamics Group, Mississauga, Ontario, Canada) at 20 mW/cm$^2$ for 60 sec. Finally, the composite complex core-shell structures were placed in PBS to evaluate maintenance of their original organized structure.

2.8.3. Characterization and chondrogenic differentiation of hMSCs encapsulated 4-OMA microgel

To confirm whether GMs were located within the core site with hMSCs or not, core/shell microgels were generated with hMSCs only or hMSCs/empty GMs encapsulated within the core of 4-OMA shells. The density of GMs was 0.1875 mg with 0.25 x 10$^6$ hMSCs.[7]

1 ml gastight syringes was then filled with each of these mixtures. Next, microgels were generated via the above described process. They were incubated for 21 days in a basal pellet medium (BPM), which consisted of HG-DMEM with 1 % ITS™ + Premix (Corning®), 100 nM dexamethasone (Sigma Aldrich), 37.5 μg/ml of l-ascorbic acid 2-phosphate (Sigma Aldrich), 1 mM sodium pyruvate (HyClone), and 100 μM nonessential amino acids (HyClone). We also cultured two core/shell microgels together to evaluate their ability of their cores to fuse together. The cellular morphology changes were evaluated at 0, 1, 3, 7, 14, and 21 days post-culture using inverted phase-contrast microscopy (TMS-F, Nikon Corporation, Tokyo, Japan) (n=4). To visualize the cellular cytoskeleton, filamentous actin (F-actin) in aggregates was stained with Alexa Fluor™ 488 Phalloidin (Cat. No. A12379, Thermo Fisher Scientific) with DAPI counterstain after 21 days of culture. Briefly, aggregates were fixed in a 3.7% paraformaldehyde solution in PBS for 1 h at room temperature followed by rinsing with PBS three times. Aggregates were then permeabilized for 20 min by 0.1% triton X-100 diluted in PBS and subsequently rinsed with PBS three times. After that, they were blocked with 1 % BSA for 30 min and subsequently rinsed with PBS three times. Finally, aggregates were stained using Alexa Fluor 488 conjugated F-actin staining solution (diluted 1:200 in PBS) for 1 h in the dark by wrapping the samples with aluminium-foil. Before capturing images, the samples were counterstained with the DAPI solution for 5 min. Fluorescence images were obtained with a fluorescence microscope (Nikon Eclipse TE300, Tokyo, Japan). To determine whether GMs also stained with Alexa Fluor™ 488 Phalloidin, unincorporated GMs were stained with Alexa Fluor™ 488 Phalloidin as described above.

To confirm the chondrogenic differentiation of hMSCs encapsulated in 4-OMA core/shell microgels, microgels were formed under various conditions and cultured in different media compositions. The detailed experimental groups and media compositions are described in Table S2 and Figure S7. Chondrogenic pellet medium (CPM) consisted of BPM supplemented with 10 ng/ml TGF-β1. The chondrogenic differentiation of hMSCs encapsulated in 4-OMA core/shell microgels was measured by glycosaminoglycan (GAG) assay after 2 and 3 weeks of culture. The hMSC aggregates were digested by papain buffer solution (25 μg/mL papain, 2 mM L-cysteine, 50 mM sodium phosphate, 2 mM ethylenediaminetetraacetic acid) in EP-Tubes at 65 °C overnight. After that, GAG and DNA content were measured using the DMMB assay at pH 1.5[3a, 8] and the PicoGreen assay,[2, 7b, 9] respectively, according to the manufacturers' instructions (n=4).

2.8.4 Histological analysis and immunofluorescence staining after *in vitro* chondrogenic differentiation at 3 weeks

To histologically evaluate chondrogenesis, safranin O staining with fast green counterstain, toluidine blue O staining, and immunofluorescence staining for collagen type II were performed. Before staining, all samples were frozen in Optimal Cutting Temperature (OCT, Fisher Scientific, 23-730-573, Pittsburgh, PA) compound and then cut with a Microm HM505E cryostat (Microm GmbH, Walldorf, Germany) at -20 °C at 14 μm in thickness. The sections were collected on a glass microscope slide and the OCT compound was removed by incubating in ultrapure deionized Milli-Q® water for 15 min. After that, the sections were carefully washed with PBS. For safranin O with fast green and toluidine blue O staining, slides were washed with water and dehydrated using a series of 70%, 95% and 100% ethanol. For safranin O with fast green staining, slides were stained with fast green solution (0.05%) for 1 min followed by rinsing with acetic acid solution (1 %, v/v) for 10 sec. Samples were then stained in 0.1% safranin O solution for 5 min. Finally, they were dehydrated with 95% and 100% EtOH and fixed using Permount™ mounting medium (UN1294, Fisher Scientific, Pittsburgh, PA). For toluidine blue O staining, 0.05 % toluidine blue O (pH 4) solution in McIlvaine buffer [Citric Acid (100mM) / sodium phosphate dibasic (200mM)] was applied to the samples for 4 min, and then they were washed. The slides for toluidine blue O staining were prepared the same as those for safranin O with fast green staining. For immunofluorescence staining against collagen type II, OCT compound was removed by immersing the slides in ultrapure deionized Milli-Q® water for 15 min. After removing excess water, the slides were then permeabilized for 20 min using 0.1% triton X-100 diluted in PBS and then rinsed with PBS three times. After that, they were blocked with 1 % BSA for 30 min and then rinsed with PBS three times. Finally, the slides were stained using rabbit polyclonal anti-collagen type II antibody (diluted 1:200 in PBS, Abcam, Cat No. ab34712, Cambridge, MA) overnight at 4°C. They were then washed carefully with PBS. Finally, they were incubated with goat anti-rabbit Alexa Fluor® 488 secondary antibody (diluted 1:200 in PBS, Cat No. 111-545-144, Jackson ImmunoResearch Laboratories, Inc., West Grove, PA) for 1 h in the dark and then washed using PBS again. Next, slides were mounted using Fluoromount™ aqueous mounting medium (Cat. No. F4680, Sigma-Aldrich). Finally, all photomicrographs of stained samples were taken using a microscope (Nikon Eclipse TE300, Tokyo, Japan).

2.9 *In vivo* analysis

For *in vivo* assessments, CB-17 female severe combined immunodeficiency (SCID) mice (5-6 weeks old for the hydrogel degradation study and 9 weeks old for the chondrogenesis study) were obtained from Taconic Biosciences (Albany, NY). All animals were treated in accordance with the National Institutes of Health's (NIH) *Guidelines for the Care and Use of Laboratory Animals* and an Institutional Animal Care and Use Committee (IACUC) of the University of Illinois at Chicago approved protocol (Protocol No.: 19-152). Before surgery, mice were anesthetized initially with a mixture of 3 % isoflurane in $O_2$ and then maintained on 1.5-2 % isoflurane in $O_2$. The flow rate of $O_2$ was 1L/min. Before incision, mice were administered buprenorphine (concentration: 0.03 mg/ml, dose: 0.1 mg/kg) under veterinarian guidance. An incision was made, and subcutaneous pockets were prepared on the backs of the mice. High-density cells encapsulated core/shell microgels were implanted into the pockets, which were then closed with interrupted suture of 5-0 nylon monofilament (Cat. No. 668G, Ethicon, Inc., Somerville, NJ). All animal procedures were carried out on a heating pad to maintain animal body temperature. At predetermined time points, the animals were sacrificed by using 100% $CO_2$ gas in a chamber.

2.9.1 Characterization of *in vivo* hydrogel degradation and tissue integration at 3 and 7 days post-implantation

To verify the degradation of the constructs' hydrogel shell *in vivo* and evaluate the resultant core tissue integration with host tissue, hMSCs encapsulated in unmodified alginate and 4-OMA core/shell microgels were implanted into left and right dorsal subcutaneous pouches of mice immediately after the constructs were made. The generation method of cell encapsulated core/shell microgels was same as described for the *in vitro* studies. The microgels were implanted bilaterally with the unmodified alginate constructs on the left dorsal side and the 4-OMA constructs on the right dorsal side. All mice exhibited no adverse reaction to the presence of the implants for the duration of the experiment. All animals survived until they were sacrificed. Implanted core/shell microgels, along with surrounding host tissues, were harvested at 3 and 7 days post-surgery, and then immediately fixed in 10% neutral-buffered formalin (NBF) for 24 h for histological analysis. After the series of ethanol, xylene-ethanol, and xylene treatment, samples were embedded in paraffin and sectioned at 7 μm thickness by rotary microtome (Leica RM2255, Leica Microsystems, Wetzlar, Germany). Sections were deparaffinized and rehydrated prior to histological staining. Sections were then stained with 1% alcian blue (pH 2.5) with nuclear fast red as a counterstain and other sections were stained with haematoxylin and eosin (H&E).

2.9.2 Characterization of *in vivo* cartilage formation and host tissue integration

For investigation of chondrogenesis and fusion of the implanted construct with host tissue *in vivo*, core/shell microgels were implanted as described above. To drive chondrogenesis *in vivo*, we implanted various groups described in Table S3 and Figure S8. Fused microgels with surrounding host tissue were harvested at 1 and 3 weeks post-surgery. For quantification of GAG/DNA content, harvested samples (n=5) at 3 weeks were analyzed by the GAG and DNA assays as described above.

During the *in vivo* chondrogenesis study, samples were harvested at 1 week post-surgery to evaluate the integration of the implanted constructs with host tissue. All mice exhibited no adverse reaction to the presence of the implants for the duration of the experiment. All of them survived until they were sacrificed. All samples underwent the same tissue processing as described above, and then histological staining was performed. H&E staining and 1% alcian blue (pH 2.5) staining with nuclear fast red as a counterstain were performed. For immunohistochemistry staining of human nuclear antigen (HNA) to distinguish between implanted hMSCs and host murine tissue, slides were processed via standard methodology. In detail, the glass slides were deparaffinized and rehydrated in the graded series of ethanol, and then they were carefully rinsed with ultrapure deionized Milli-Q® water. Next, sodium citrate buffer (10 mM sodium citrate dihydrate, 0.05% Tween 20, pH 6.0) was placed in a slide container rack and heated in a vegetable steamer to 98 °C, and then the glass slides were placed in pre-heated sodium citrate buffer in the slide container rack. The antigen retrieval process was performed for 20 min at 98 °C, and then the glass slides in the slide container rack were cooled down to room temperature for 30 min on a laboratory bench. After washing with PBST (PBS + 0.1% Tween 20 detergent) two times for 5 min, endogenous peroxidases were blocked with 3% hydrogen peroxide diluted in PBS (Reagent ACS, Cat. No. 470301-282, Ward's

Science, Rochester, NY) for 30 min. After washing with PBST two times, slides were carefully incubated with 2.5% Normal Horse Serum (ImmPRESS kit, Vector Laboratories, Burlingame, CA) to block non-specific binding for 2 h. After removal of excess serum from sections without a washing step, the slides were incubated with primary mouse anti-Human IgG1 Nuclei (Cat. No. MAB1281, Millipore, CA, 1:100 diluted in 1 % normal horse serum) overnight at 4°C in a humidified chamber. After carefully washing the slides with PBST, they were incubated in ImmPRESS™ HRP Anti-mouse IgG (Peroxidase) Polymer Detection Kit derived from horse serum (Cat. No. MP-7402, Vector Laboratories, Burlingame, CA) for 30 min. Next, slides were processed with ImmPACT™ DAB Eqv Substrate Kit (Cat. No. SK-4103, Vector Laboratories, Burlingame, CA) for visualization. After washing with ultrapure deionized Milli-Q® water, the slides were stained by Mayer's hematoxylin (Cat. No. TA-125-MH, Lab Vision, Fremont) as a counter stain. For control staining, host only tissue (group 3 constructs) was stained using the same staining method as above and a group 3 condition slide was stained using an IgG1 isotype control from murine myeloma (Cat. No. M5284, Sigma-Aldrich, St. Louis, MO) instead of the HNA primary antibody. Finally, all slides were mounted using Permount™ mounting medium (UN1294, Fisher Scientific, Pittsburgh, PA). Then, all slides were imaged using a microscope (Nikon Eclipse TE300, Tokyo, Japan).

At the 3 week harvest, H&E staining, safranin O with fast green, toluidine blue O (0.05%, pH 4.0), aldehyde fuchsin (pH 1, Cat. No. 26328-01, Electron Microscopy Sciences, Hatfield, PA) with 1% alcian blue in 0.1 M HCl (pH 1), Picrosirius red (Cat. No. 24901–500, Polysciences Inc., Warrington, PA), and immunohistochemistry against HNA were performed to visualize tissue morphology, sulfated GAG content, and generated cartilage by the implants. H&E, safranin O with fast green, toluidine blue O, and immunohistochemistry against HNA were performed as described above. Picrosirius red staining was conducted following the manufacturer's instructions. For aldehyde fuchsin with alcian blue staining,[10] deparaffinized and hydrated slides were rinsed with 0.1 N HCl (pH 1). Then, the slides were stained by aldehyde fuchsin solution (pH 1) for 30 min. After rinsing with 0.1 N HCl, the slides were stained with a 1% alcian blue solution (pH 1). After rinsing again, all slides were mounted using Permount™ mounting medium (UN1294, Fisher Scientific, Pittsburgh, PA) and then imaged on a microscope (Nikon Eclipse TE300, Tokyo, Japan).

2.10 Statistical analysis

All values are depicted as mean ± standard deviation. Statistical analysis of the diameter of tissue aggregates (Figure S4d) and quantified DNA content normalized to day 0 (Figure S5b) was performed using two-way analysis of variance (ANOVA) with Bonferroni *post-hoc* test in GraphPad® Prism 5.0 (GraphPad Software, San Diego, CA). Comparison was performed against all groups in these experiments, but no significant differences were found ($p > 0.05$). In addition, statistical analysis of GAG, DNA, and GAG/DNA content after *in vitro* chondrogenic differentiation (Figure 4a) and *in vivo* chondrogenic differentiation (Figure 7b) was performed the same was as described above. Values of $p^* < 0.05$, $p^{**} < 0.01$, and $p^{***} < 0.001$ were considered statistically significant. No statistically significantly differences were presented as "N / D".

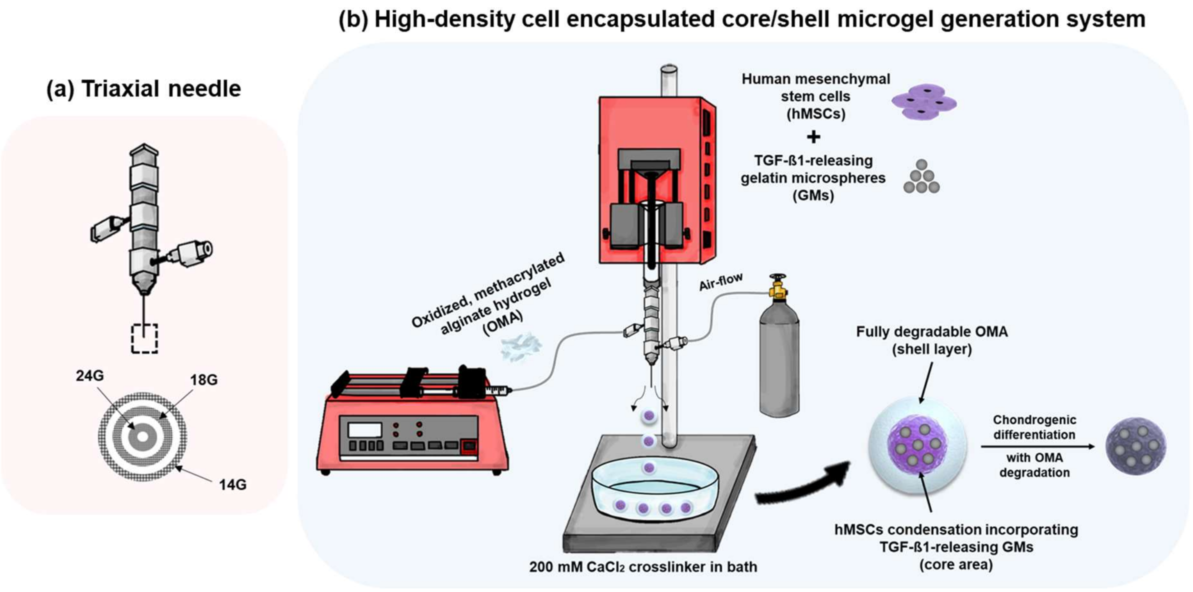


**Figure S1. Detailed schematic of immediately implantable high-density hMSCs with TGF-β1-releasing GMs encapsulated core/shell OMA microgel carrier.** Illustration of (a) triaxial nozzle structure and (b) generation method via triaxial air-flowing system. hMSCs & GM core/ hydrogel shell co-axial flow is initially formed, and then airflow in the most exterior layer separates core-shell droplets in a spherical particle shape. These droplets are collected in a bath containing a $CaCl_2$ solution for direct crosslinking of the outer macromer solution to form a hydrogel shell. In this study, the core-shell microgels were cultured in vitro or immediately implanted subcutaneously in mice. Degradable OMA shell layers degrade within several days during culture leaving a cell condensation. TGF-β1 in the cell condensation drives chondrogenic differentiation of the hMSCs and formation of cartilage-like tissue, and the resulting aggregate tissues can fuse together with each other and surrounding host tissue.

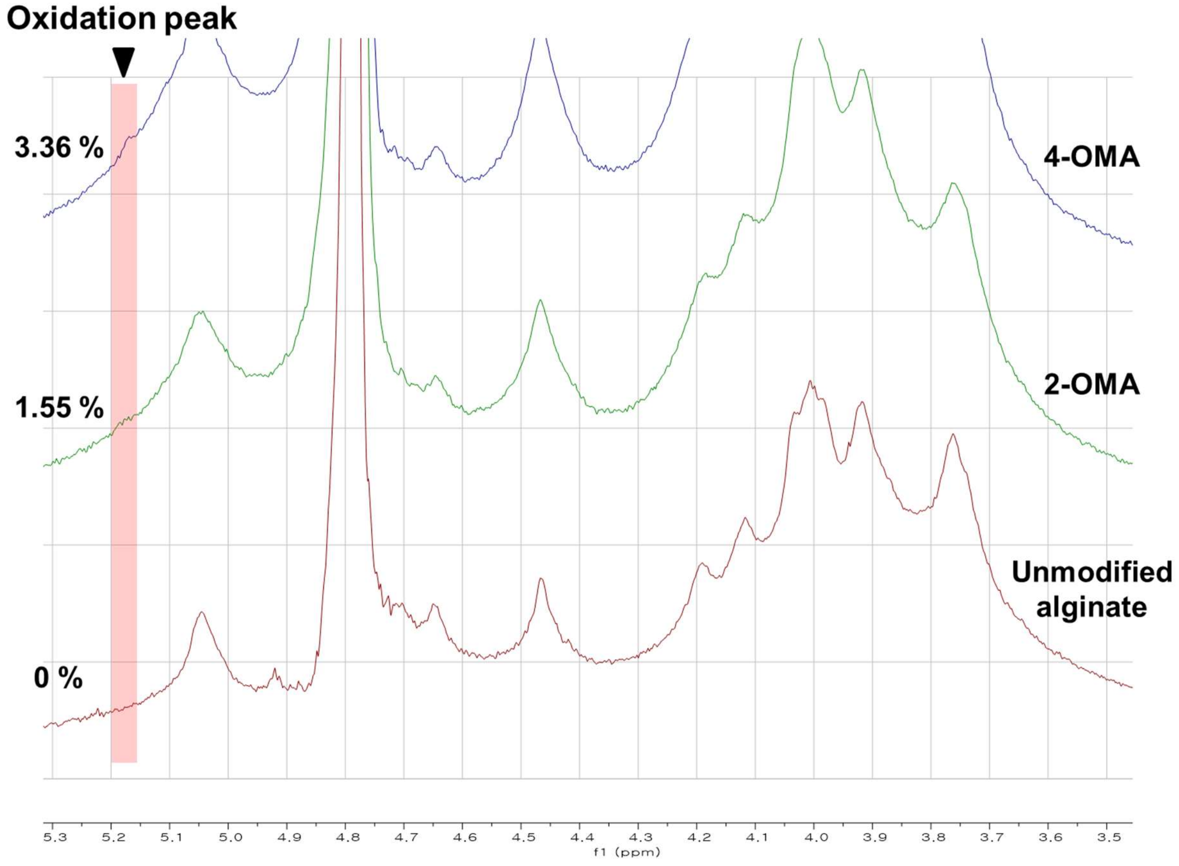


**Figure S2. Quantification of oxidation degree of OA hydrogels via [1]H-NMR analysis.** The actual oxidation degrees (theoretical: 2 and 4 %) of oxidized alginates were 1.55 and 3.36 %, respectively. The oxidation degree was increased by increasing the amount of sodium periodate used during OMA synthesis.

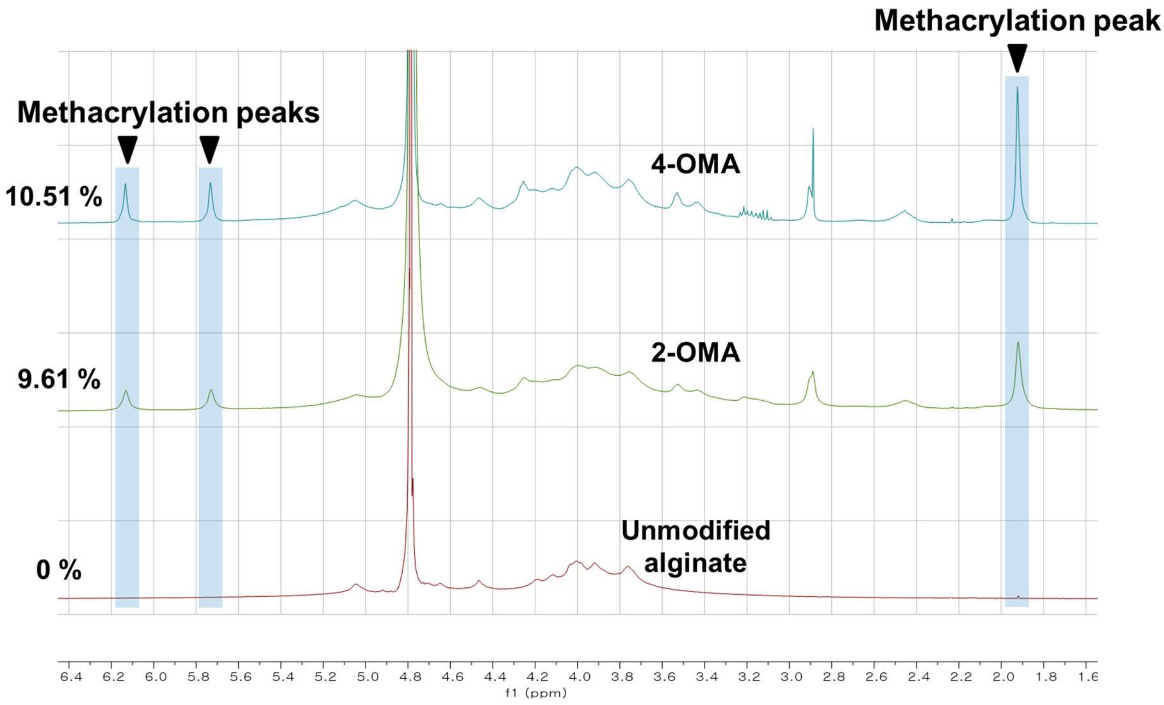


**Figure S3. Quantification of methacrylation degree of OMA hydrogels via $^1$H-NMR analysis.** The actual methacrylation degrees of 2-OMA and 4-OMA (theoretical: 40%) were 9.61 and 10.51 %, respectively.

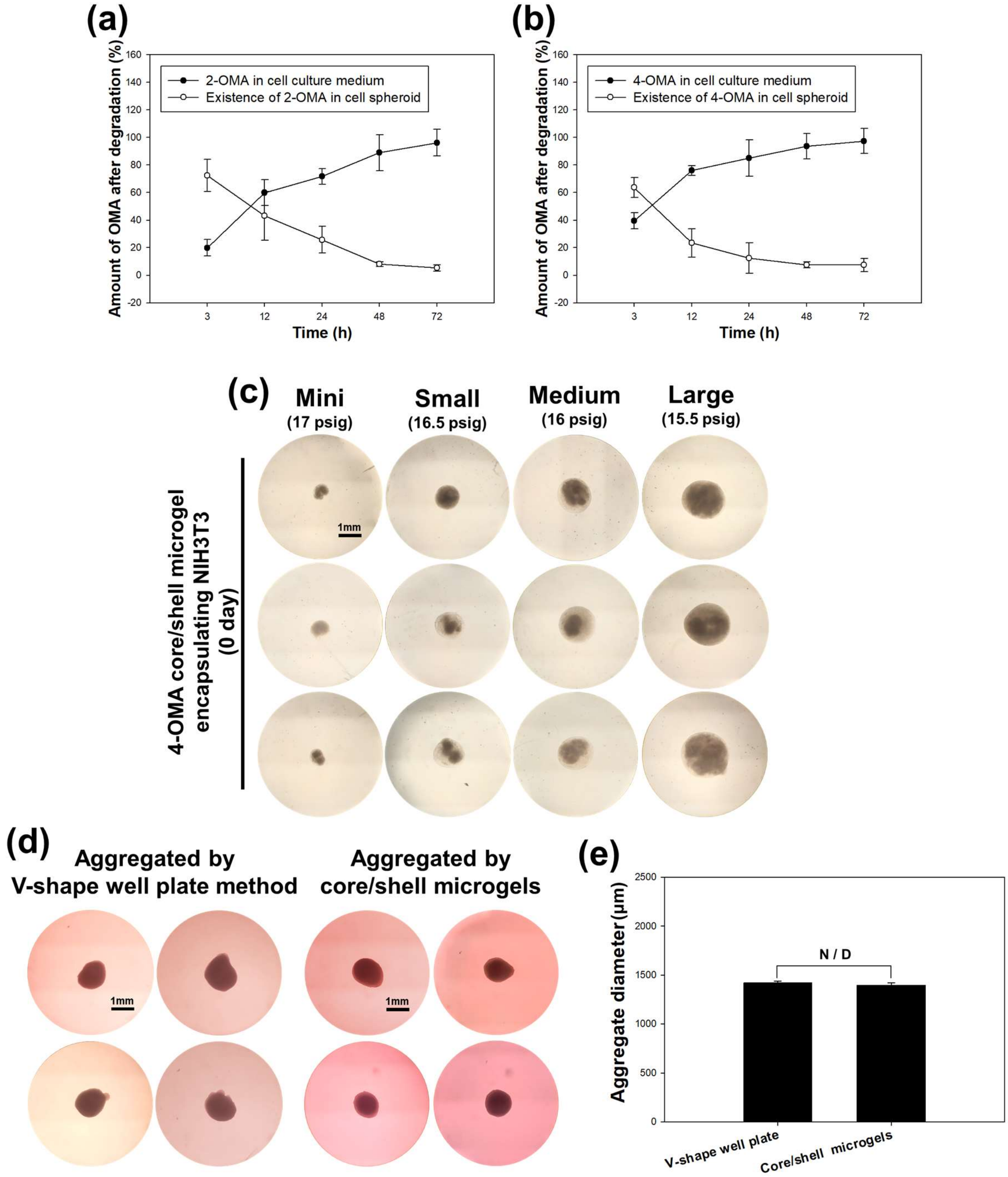


**Figure S4.** ***In vitro* degradation kinetics of NIH3T3 core/ OMA shell microgels.** (a and b) The supernatant was carefully harvested at a predetermined time point, core/shell microgels were broken down with a homogenizer, and then the amounts of degraded and residual OMA were quantified by DMMB assay. Absorbance intensities of the samples were measured at predetermined time points. **Comparison of traditional NIH3T3 cell condensations created**

**by the V-shape well plate method and (c) core / 4-OMA shell microgels.** For comparison with traditionally made 3D cell spheroids, NIH3T3 condensations were generated by the V-shape well plate method and then cultured for 7 days. Condensations formed by both methods have similar (d) 3D spheroidal morphologies and (e) aggregate diameter.

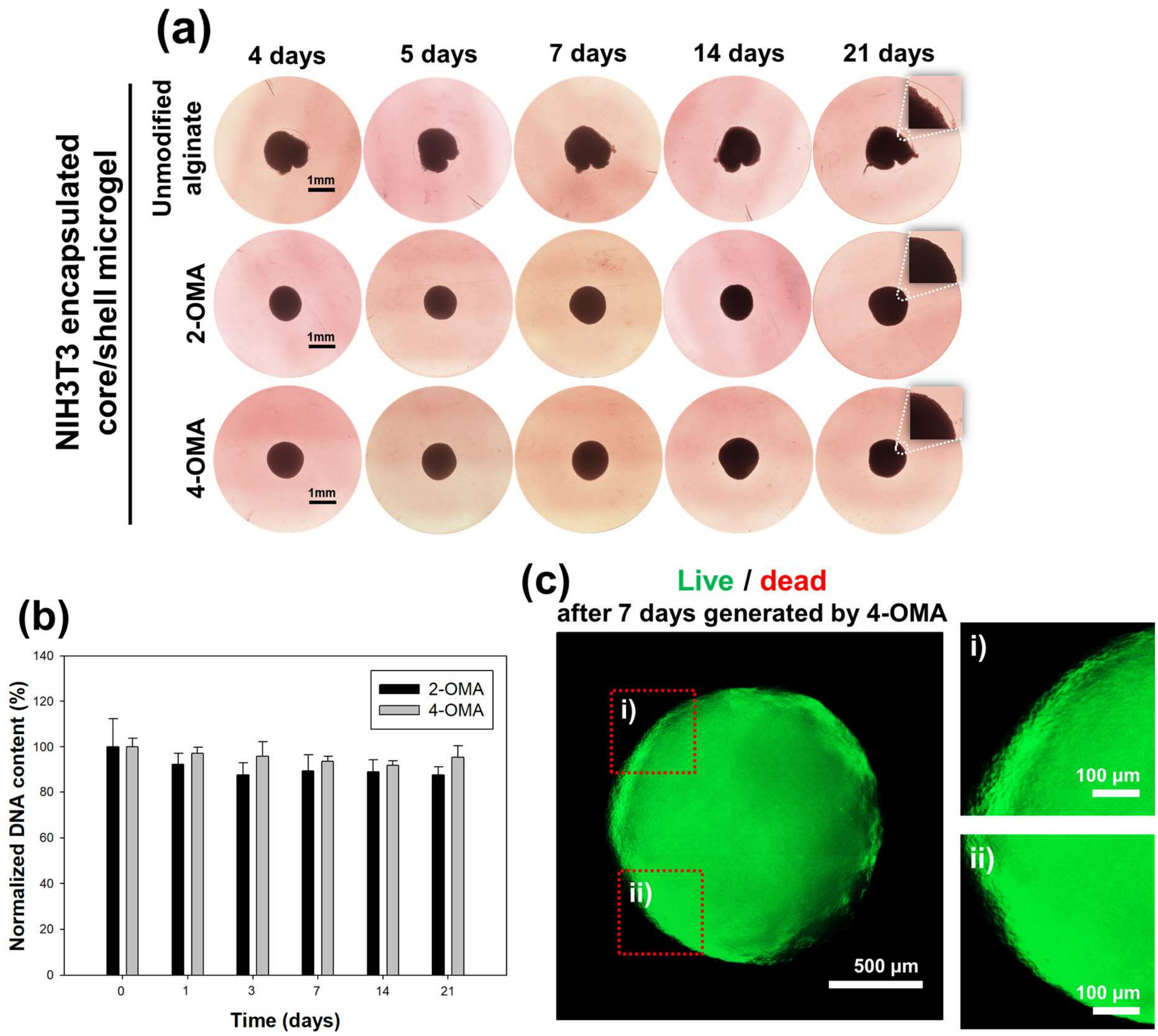


**Figure S5. Morphology change of NIH3T3 encapsulated core/shell microgels from 4 days to 21 days.** (a) In the unmodified alginate group, the hydrogel shell layer continuously maintained NIH3T3 encapsulation over 21 days of cultivation. In the 2-/4-OMA groups, NIH3T3 condensation robustly maintained round aggregate structures long after OMA shell degradation. **Quantification of total DNA content and live/dead assay**. (b) NIH3T3 core / 2-/4-OMA shell microgels were cultured for 21 days to measure DNA content. The DNA content minimally decreased over 21 days of cultivation, with no significant differences between time points ($p > 0.05$). (c) Live/dead assay was performed after 7 days of culture. Spherical living-3D cell condensations were observed as evidenced by the strong green fluorescence. High magnification images (c-i and ii) showed high-density cell condensations with tight cellular interconnection.

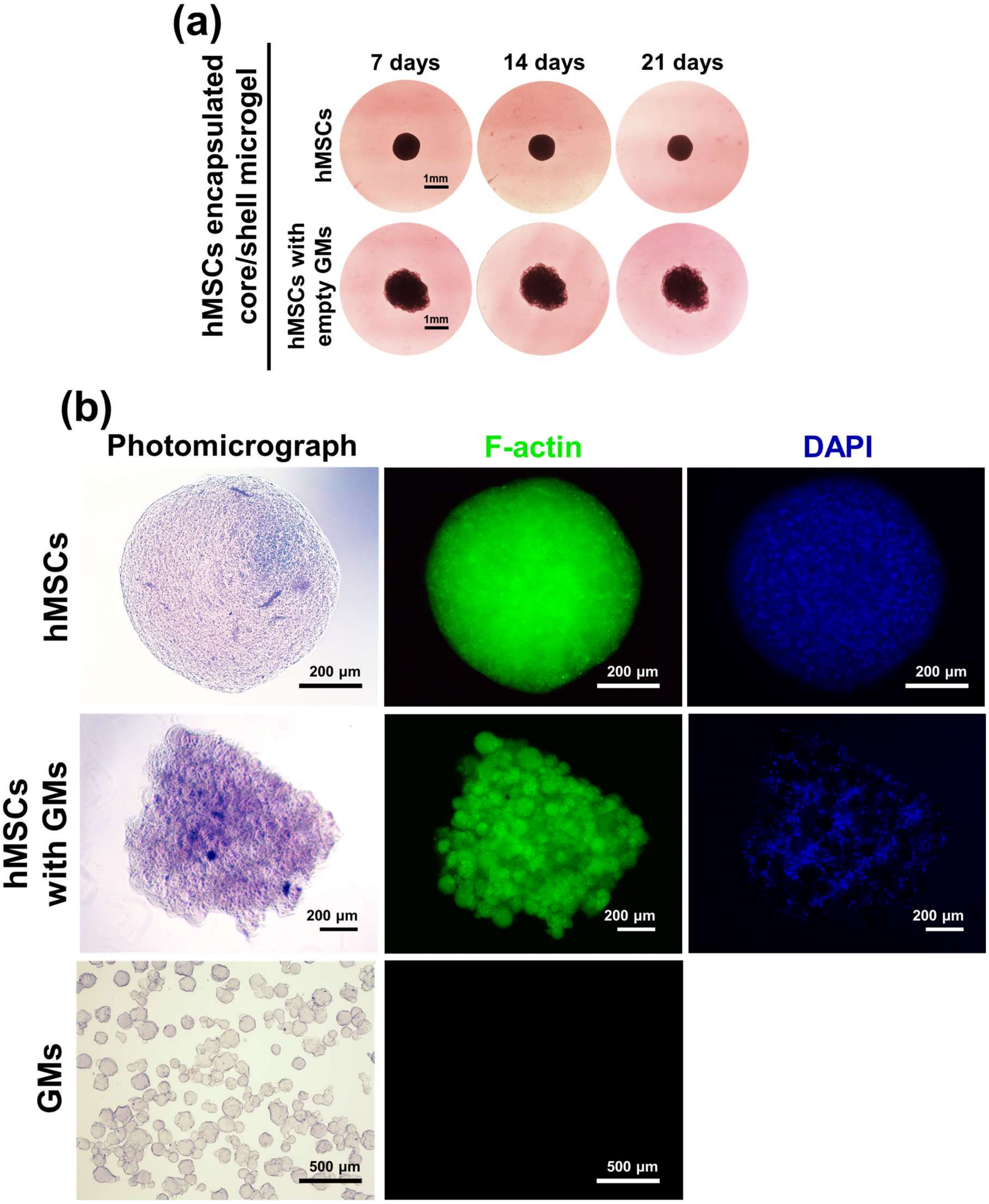


**Figure S6 Morphology change of hMSCs and hMSCs/GMs core / 4-OMA shell microgels from 7 days to 21 days.** (a) The condensation morphology did not change over 21 days of culture. **F-actin and nuclear staining of the hMSC condensations generated by 4-OMA microgel shells and visualized by Alexa Fluor™ 488 Phalloidin with DAPI staining, respectively.** (b) The hMSC and hMSC/GMs aggregates formed from core/shell microgels were processed after 21 days of culture and exhibited compact high-density cellular condensations.

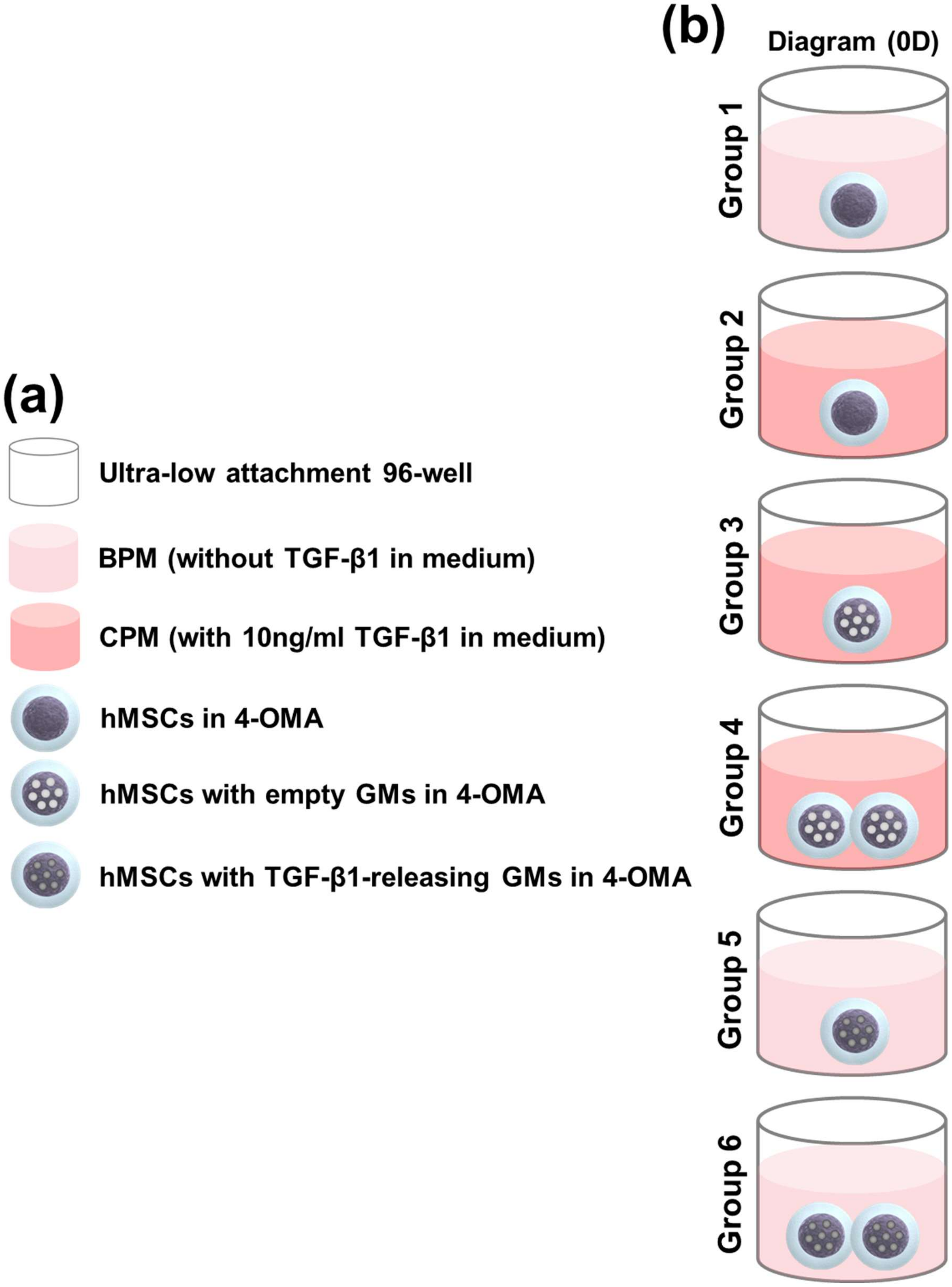


**Figure S7. A detailed schematic illustration of *in vitro* chondrogenic differentiation groups and their culture conditions.** (a) Various media compositions and experimental groups were designed and then cultured in ultra-low attachment 96-well plates (b). *In vitro*

Groups 1 had no TGF-β1 supplementation, but Group 2 had TGF-β1 supplementation in the media. Groups 3 and 4 encapsulated empty GMs but exogenous TGF-β1 was supplied in the media during culture. In groups 5 and 6, TGF-β1-releasing GMs were encapsulated in hMSC-laden microgels, but TGF-β1-free media was supplied during culture.

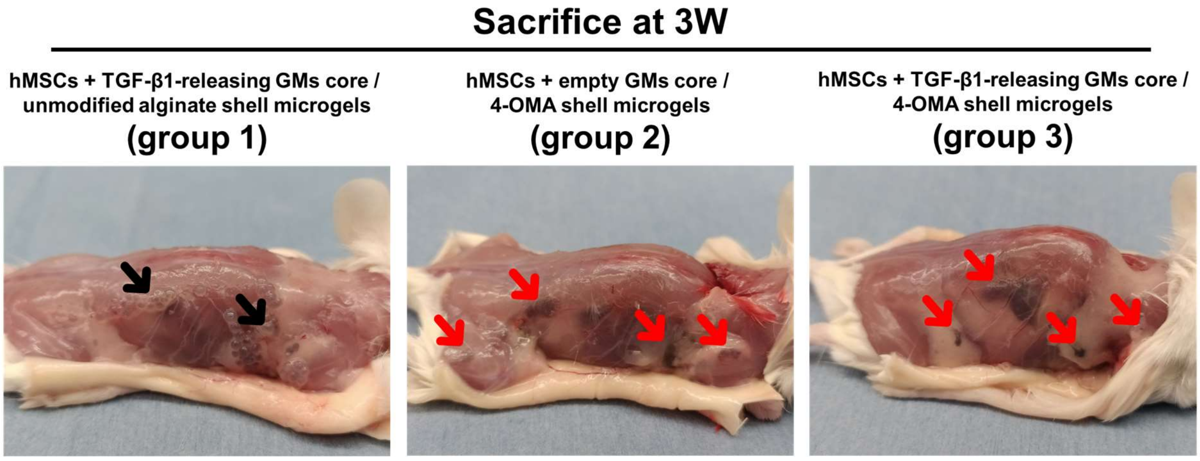


**Figure S8. Photographs of explanted constructs at 3 weeks post-surgery**. Group 1 unmodified alginate shell microgels displayed a round shape of with considerable volume. The round spheroidal shape was not seen in either group 2 or 3, indicating complete tissue fusion and integration of hMSC condensations with each other and host tissue (black arrows: undegraded microgel; red arrows: fused cell condensations after alginate shell degradation).

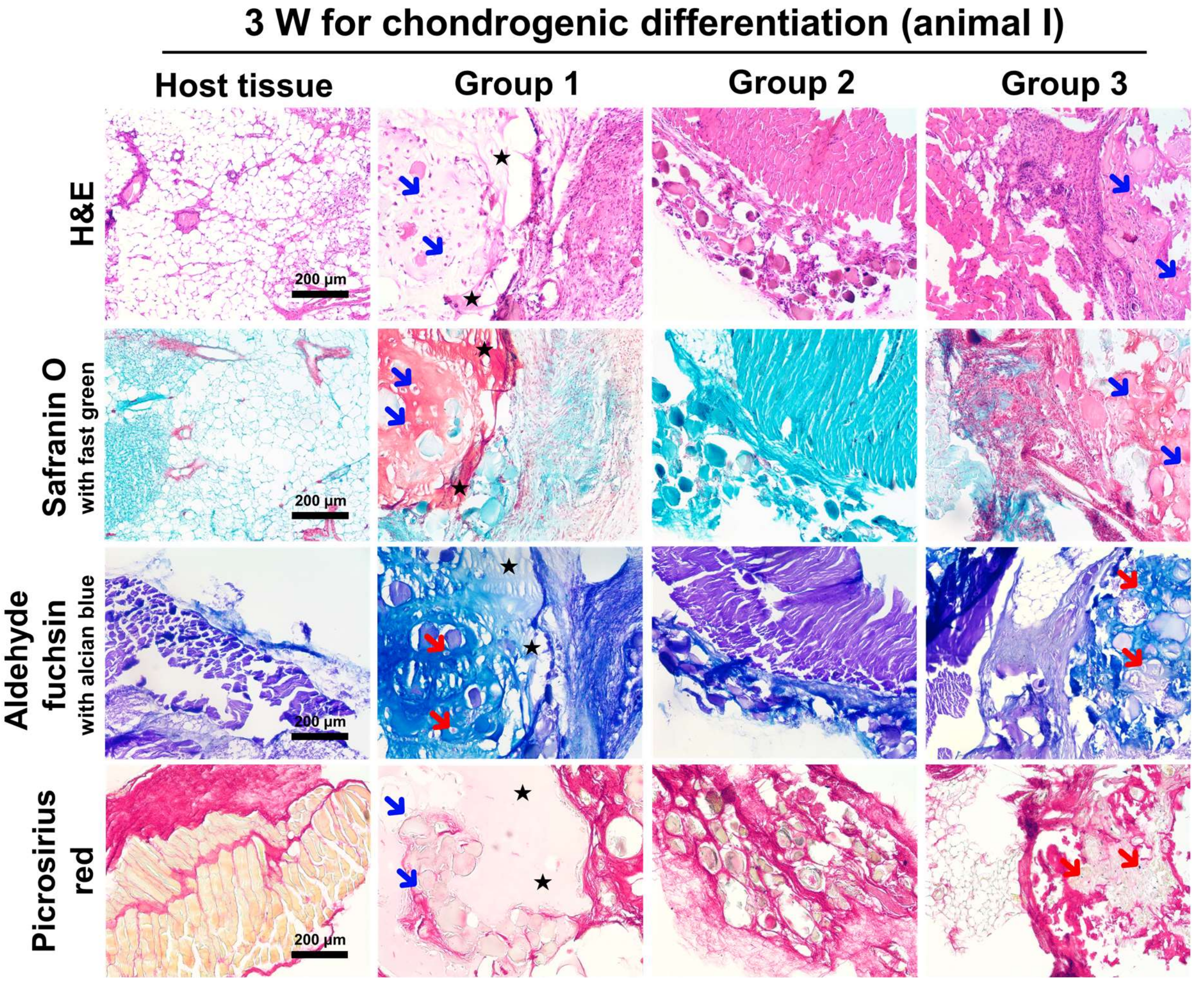


**Figure S9. Subcutaneous *in vivo* characterization of cartilage formation with its fusion depending on hydrogel types and TGF-β1-releasing GMs at 3 weeks post-surgery (animal I).** Subcutaneous cartilage tissue formation in mice was visualized by H&E, safranin O with fast green red, aldehyde fuchsin with alcian blue, and picrosirius red. In group 1, formation of cartilage matrix was good but unmodified alginate remained that interfered with tissue fusion. Group 2 showed great tissue integration but there was no cartilage matrix. In group 3, both tissue fusion and cartilage tissue formation were excellent (white and black stars; residual unmodified alginate, blue and red arrows; chondrocyte within lacunae embedded in hyaline cartilage matrix).

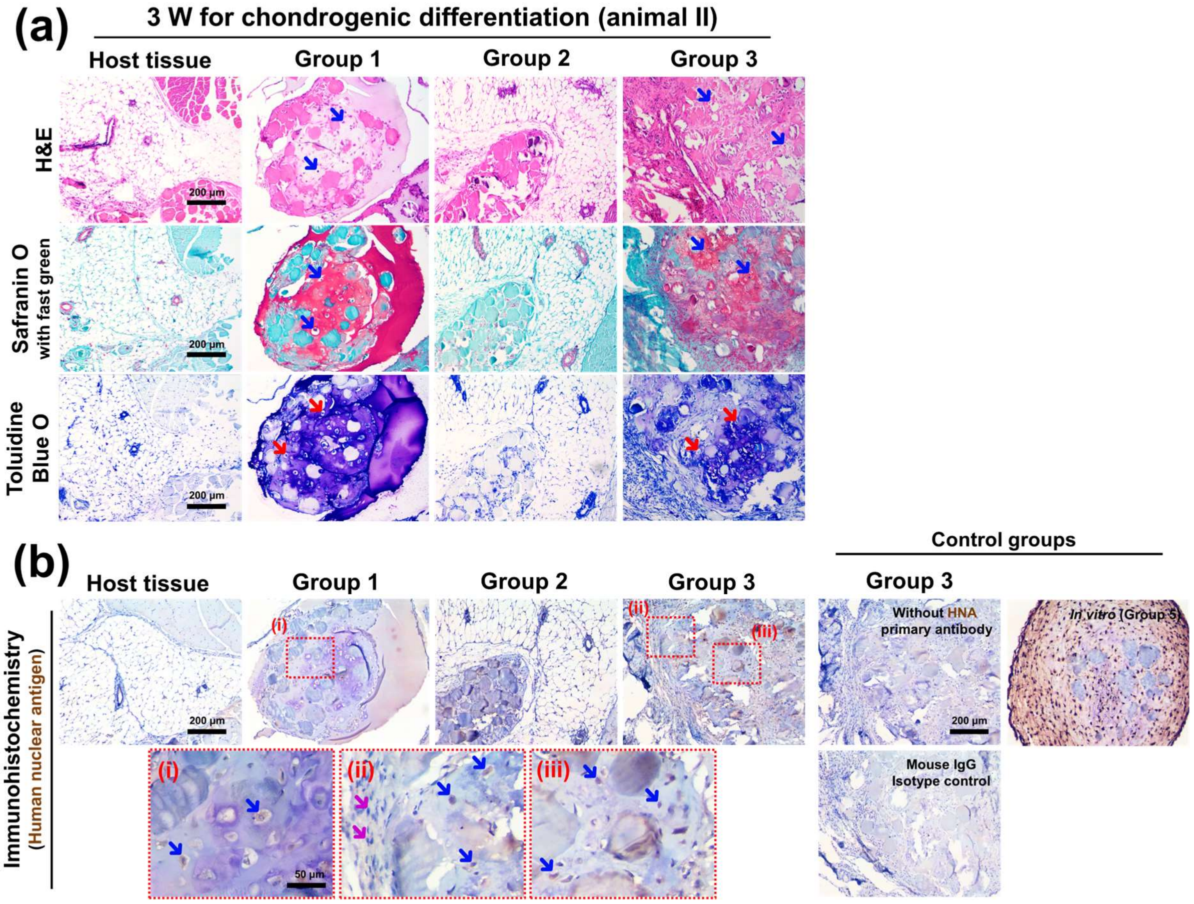


**Figure S10. Characterization of *in vivo* subcutaneous chondrogenic differentiation of hMSC/GMs encapsulated core / 4-OMA shell microgels at 3 weeks post-surgery (animal II).** (a) Subcutaneous cartilage tissue formation in mice was visualized by photomicrographs of H&E, safranin O with fast green, and toluidine blue O stained sections. Histology staining results were similar to that of animal I. (b) The hMSC-derived chondrocytes were stained by HNA with Mayer's hematoxylin as a counterstain. (b-ii) Host mouse cells stained blue in color by Mayer's haematoxylin counterstaining. However, (b-i and iii) hMSC generated chondrocytes stained brown in color, which was confirmed by control groups. Results indicated that cartilage-like matrix containing hMSC-derived chondrocytes fused with surrounding host tissue (blue and red arrows: chondrocyte within cartilage-like matrix; purple arrows: host mouse cell).

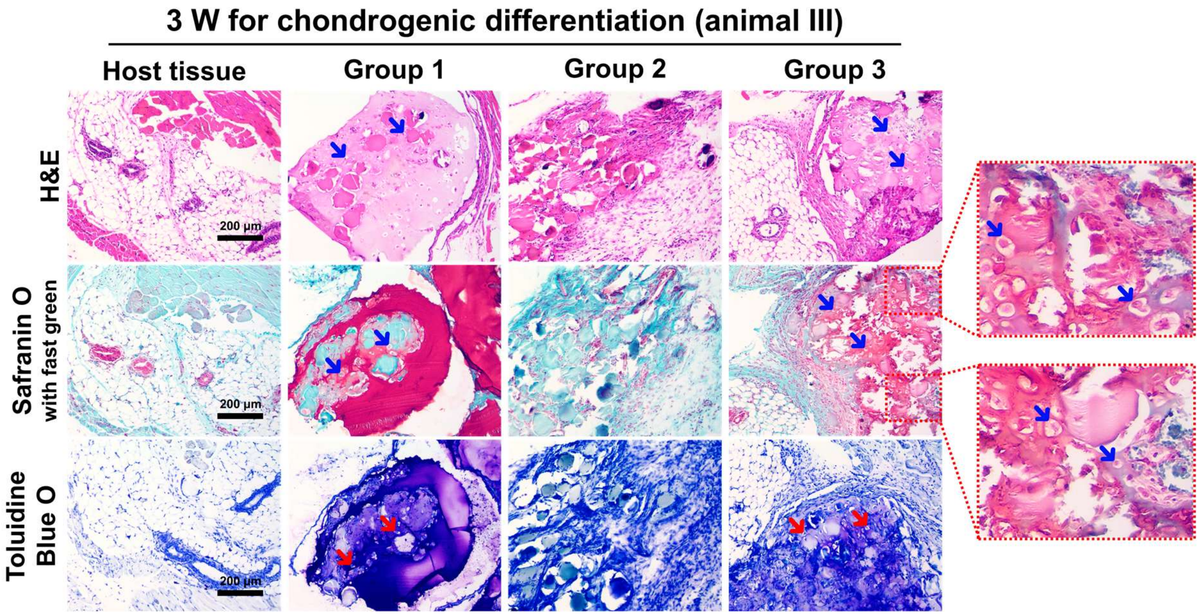


**Figure S11. Characterization of *in vivo* subcutaneous chondrogenic differentiation of hMSC/GMs encapsulated core / 4-OMA shell microgels at 3 weeks post-surgery (animal III).** Subcutaneous cartilage tissue formation in mice was visualized by (a) H&E, safranin O with fast green, toluidine blue O. Histology staining results were similar to that of animals I and II (blue and red arrows: chondrocyte within cartilage-like matrix).

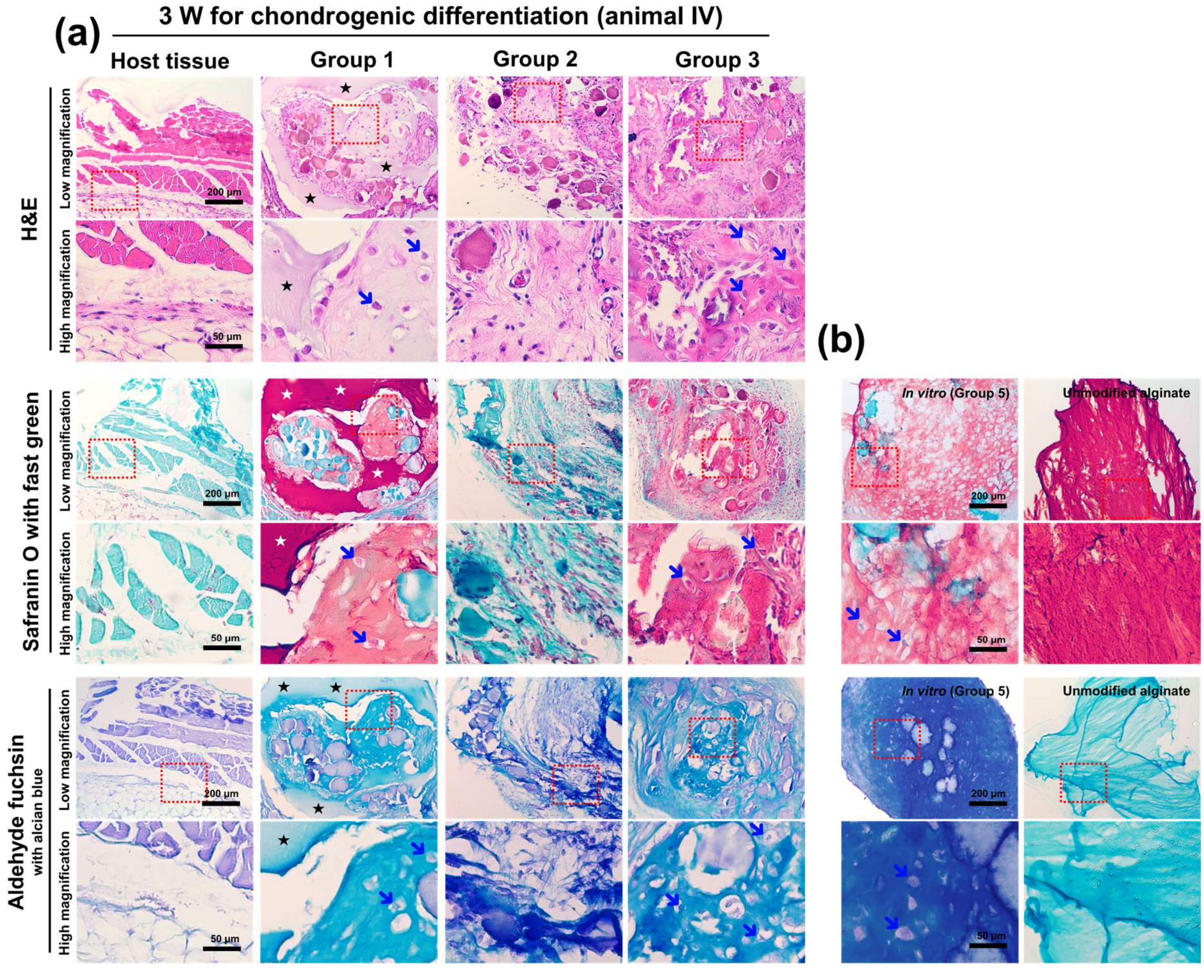


**Figure S12. Characterization of *in vivo* subcutaneous chondrogenic differentiation of hMSC/GMs encapsulated core / 4-OMA shell microgels at 3 weeks post-surgery (animal IV).** (a) Subcutaneous cartilage tissue formation in mice and (b) control groups (*in vitro* Group 5 samples, unmodified alginate without cells) were stained with H&E, safranin O with fast green, and/or aldehyde fuchsin with alcian blue and visualized through photomicrographs. Staining results were similar to those of animals I, II, and III (black and white stars: residual unmodified alginate; blue arrows: chondrocyte within cartilage-like matrix).

**Table S1.** Different types of alginate used for core/shell microgel generation

| Groups | Abbreviations | Actual oxidation and methacrylation degree (% / %) |
|---|---|---|
| 1. Unmodified alginate | N / A | 0 / 0 |
| 2. 2 % oxidized and 40 % methacrylated alginate | 2-OMA | 1.55 / 9.61 |
| 3. 4 % oxidized and 40 % methacrylated alginate | 4-OMA | 3.36 / 10.51 |

**Table S2.** *In vitro* chondrogenesis experimental groups of hMSC core / 4-OMA shell microgels

| Groups | Medium conditions |
|---|---|
| 1. hMSC incorporated core/shell microgel (single) | BPM (without TGF-β1) |
| 2. hMSC incorporated core/shell microgel (single) | CPM (BPM + 10 ng/ml of TGF-β1) |
| 3. hMSC + empty GMs core/shell microgel (single) | CPM (BPM + 10 ng/ml of TGF-β1) |
| 4. hMSC + TGF-β1-releasing GMs core/shell microgel (double) | CPM (BPM + 10 ng/ml of TGF-β1) |
| 5. hMSC + TGF-β1-releasing GMs core/shell microgel (single) | BPM (without TGF-β1) |
| 6. hMSC + TGF-β1-releasing GMs core/shell microgel (double) | BPM (without TGF-β1) |

**Table S3.** *In vivo* experimental conditions of hMSC incorporated core/shell microgels for subcutaneous implantation in mice

| Groups | Quantity (ea / pouch) |
|---|---|
| 1. hMSCs + TGF-β1-releasing GMs core / unmodified alginate shell microgels | 30 |
| 2. hMSCs + empty GMs core / 4-OMA shell microgels | 30 |
| 3. hMSCs + TGF-β1-releasing GMs core / 4-OMA shell microgels | 30 |
